# Interleukin 31 receptor alpha induces airway hyperresponsiveness in asthma

**DOI:** 10.1101/2022.12.15.520615

**Authors:** Santoshi Akkenepally, Dan JK Yombo, Sanjana Yerubandi, Bhanuprakash R. Geereddy, Francis X. McCormack, Satish K Madala

## Abstract

Asthma is a chronic inflammatory airway disease characterized by airway hyperresponsiveness (AHR), inflammation, and goblet cell hyperplasia. Both Th1 and Th2 cytokines, including IFN-γ, IL-4, and IL-13 have been shown to induce asthma; however, the underlying mechanisms remain unclear. We observed a significant increase in the expression of IL-31RA, but not its cognate ligand IL-31 during house dust mite- and *Schistosoma mansoni* soluble egg antigen-induced allergic asthma. In support of this, IFN-γ and Th2 cytokines, IL-4 and IL-13, upregulated IL-31RA but not IL-31 in airway smooth muscle cells (ASMC). Importantly, the loss of IL-31RA attenuated AHR but had no effects on inflammation and goblet cell hyperplasia in allergic asthma or mice treated with IL-13 or IFNγ. Mechanistically, we demonstrate that IL-31RA functions as a positive regulator of muscarinic acetylcholine receptor 3 (CHRM3) expression and calcium signaling involved in the contractility of ASMC. Together, these results identified a novel role for IL-31RA in ASMC contractility and AHR distinct from airway inflammation and goblet cell hyperplasia in asthma.

**Summary:** The study identified an important role for the IL-31RA-CHRM3 axis in inducing airway hyperresponsiveness with limited changes in inflammation in allergic asthma. IL-31RA, whose expression is regulated by both Th1 and Th2 cytokines, augments the CHRM3-dependent contractility of ASMC and AHR.

## INTRODUCTION

Asthma is a common chronic inflammatory disease that affects more than 300 million people worldwide. Asthma is associated with significant health and economic burdens, affecting the quality of life and decreasing productivity (*1, 2*). Despite advances in understanding the disease physiopathology and the development of anti-inflammatory therapeutic interventions, the incidence of asthma, including uncontrolled and severe forms, continues to increase. The disease is characterized by episodes of airflow obstruction that are associated with high mortality when severe and unreversed (*3*). The primary clinical manifestations in the pathophysiology of chronic asthma include airway hyperresponsiveness (AHR), inflammation, goblet cell hyperplasia with mucus production, and fibrotic airway remodeling (*4*). AHR or the excessive response of the airways to a variety of stimuli is one of the cardinal features of asthma, but AHR can also be present in other airway disorders such as cystic fibrosis and COPD(*5, 6*). Several published studies have implicated excessive airway smooth muscle contraction, airway inflammation, dysregulated neural control, and fibrotic airway remodeling as key mechanisms underlying AHR. Therefore, identifying a common pathway through which a number of these potential mechanisms operate could facilitate the development of novel therapies against AHR in asthma and other chronic lung diseases.

Traditionally, asthma was classified based on definable phenotypes, often based on features such as a clearly identifiable allergic component; however, more recently, (*7*) “endotypes” delineated by cellular and molecular pathophysiologic mechanisms that reflect mechanistic heterogeneity have been established. Currently, they include immune response endotypes driven not only by Th2 but also by Th1 (IFNγ) or Th17 (IL-17) in case of severe forms of asthma which are often associated with eosinophilia and neutrophilia (8–13). Th2 cytokines, IL-4 and IL-13, reported to be elevated in the lungs and serum of asthmatic patients, are now well-established drivers of AHR and airway inflammation in asthma (*14–16*). IL-13 is a major cytokine responsible for goblet cell hyperplasia, tissue remodeling, and bronchospasm. Even in the absence of an allergen exposure or inflammation, IL-13 alone is sufficient to induce AHR and airway inflammation in mice (*17, 18*). IL-4 is crucial in the regulation of Th2 cell proliferation as well as the antibody class switch to produce IgE, and recent studies highlight a shared role of this cytokine with IL-13 in the induction of AHR and inflammation through their common receptor type II IL-4R, as well as through type I IL4R (*19, 20*). Clinical trials that neutralize the common IL-4R receptor for these Th2 cytokines have shown improvements in AHR and inflammation in both allergic asthma and atopic dermatitis patients (*21, 22*). However, these therapies and other more conventional treatments for asthma have not been effective against low type 2 or non-type 2 endotypes and severe asthma with a mixed Th1/Th2 cytokine phenotype (*23*). However, understanding the mechanisms that regulate AHR and therapeutic resistance in these specific conditions may help to develop alternative and effective therapeutic approaches to control asthma.

IL-31 is a Th2-related cytokine implicated in tissue remodeling in chronic lung and skin diseases (*24, 25*), including allergic asthma, atopic dermatitis (AD), fibrosis, and itch-associated conditions (*24–28*). IL-31 specifically binds to a heterodimeric receptor complex formed by the IL-31 receptor alpha (IL-31RA) and OSMRβ and signals through MAPK, JAK, STAT1, STAT3, and STAT5 (*29, 30*). IL-31 is predominantly produced by hematopoietic cells, whereas its receptor, IL-31RA, is predominantly expressed in non-hematopoietic epithelial and smooth muscle cells (*31–34*). Although studies have reported an association between the expression of both IL-31 and IL-31RA with severe asthma (*35, 36*), the focus of most prior studies has been on tissue remodeling observed in diseases such as pulmonary fibrosis. For example, we recently published a bleomycin model of pulmonary fibrosis in which the absence of IL-31RA signaling had a protective effect against the impairment of lung function in fibrotic mice (*37*). In contrast, our and other previous studies have shown that Th2 cytokines, IL-4 and IL-13, augment the expression of IL-31RA in immune cells, including macrophages (*21, 38*). However, the role of the IL-31-IL-31RA axis in the induction of AHR or airway inflammation during allergic asthma remains unexplored.

In this study, we evaluated the effects of the IL-31-IL-31RA axis on allergen-induced AHR, inflammation, and goblet cell hyperplasia in mice with wild-type and IL-31RA deficient gene expression. We demonstrated a significant increase in the expression of IL-31RA, but not IL-31, and both Th1 and Th2 cytokines (IFN-γ, IL-4, and IL-13) in two alternative mouse models of allergic asthma. The loss of IL-31RA was sufficient to attenuate both allergen and cytokine (IFN-γ, and IL-13)-induced AHR but had limited or no effect on inflammation and goblet cell hyperplasia. We identified a novel mechanism underlying IL-31RA-driven AHR involving heightened muscarinic acetylcholine receptor 3 (CHRM3) expression and calcium signaling that together induces ASMC contraction. The discovery of the IL-31RA-CHRM3 axis augments calcium signaling and ASMC contraction is important and may prove beneficial in designing therapies with greater efficacy to target AHR downstream of both Th1/Th2 cytokines in asthma.

## RESULTS

### Upregulation of IL-31RA promotes airway hyperresponsiveness

Loss of type II IL-4 receptor signaling attenuates the pathological features of asthma and expression of IL-31RA during *Schistosoma mansoni* soluble egg antigen (SEA)-induced allergic asthma (*11, 20, 38-40*). We hypothesized that in a house dust mite (HDM)-induced model of allergic asthma, the loss of IL-31RA would block the development of some, if not all, pathological features of allergic asthma. To elucidate the role of IL-31RA in allergic asthma, IL-31RA knockout and wild-type mice were intraperitoneally (i.p.) sensitized twice with HDM followed by intratracheal (i.t.) challenge with HDM a week after the second sensitization for two consecutive days (Fig 1A). The control mice were sensitized and challenged with saline. As expected, the expression of IL-31RA was not detected in IL-31RA knockout (IL-31RA^-/-^) mice (Fig 1B). Sensitization and challenge with HDM or saline did not modify the gene expression of IL-31 in either wild-type or IL-31RA^-/-^ mice (Fig 1C), but significantly elevated IL-31RA in wild-type mice compared to saline-treated mice (Fig 1B). However, since HDM without alum induces allergic asthma, we assessed whether the deletion of IL-31RA alters AHR during HDM-induced allergic asthma. The loss of IL-31RA was sufficient to attenuate dose-dependent methacholine (MCh)-induced AHR in IL-31RA^-/-^ mice compared to wild-type mice sensitized and challenged with HDM (Fig 1D). To assess whether the observed decrease in AHR was due to impairment of airway contraction, we prepared precision-cut lung slices (PCLS) from wild-type and IL-31RA knockout mice and tested dose-dependent MCh-induced airway contraction in real time. MCh-induced airway contraction was decreased in PCLS prepared from the lungs of IL-31RA^-/-^ mice compared to that in wild-type mice (Fig 1E and 1F). Furthermore, we used a collagen gel contraction assay to measure carbachol-induced contraction of airway smooth muscle cells (ASMC) isolated from the trachea of wild-type and IL-31RA^-/-^ mice. Collagen gels embedded with wild-type ASMC showed higher contraction compared to IL-31RA^-/-^ ASMC within 30 minutes in floating collagen gels (Fig 1G).

**Figure 1.**
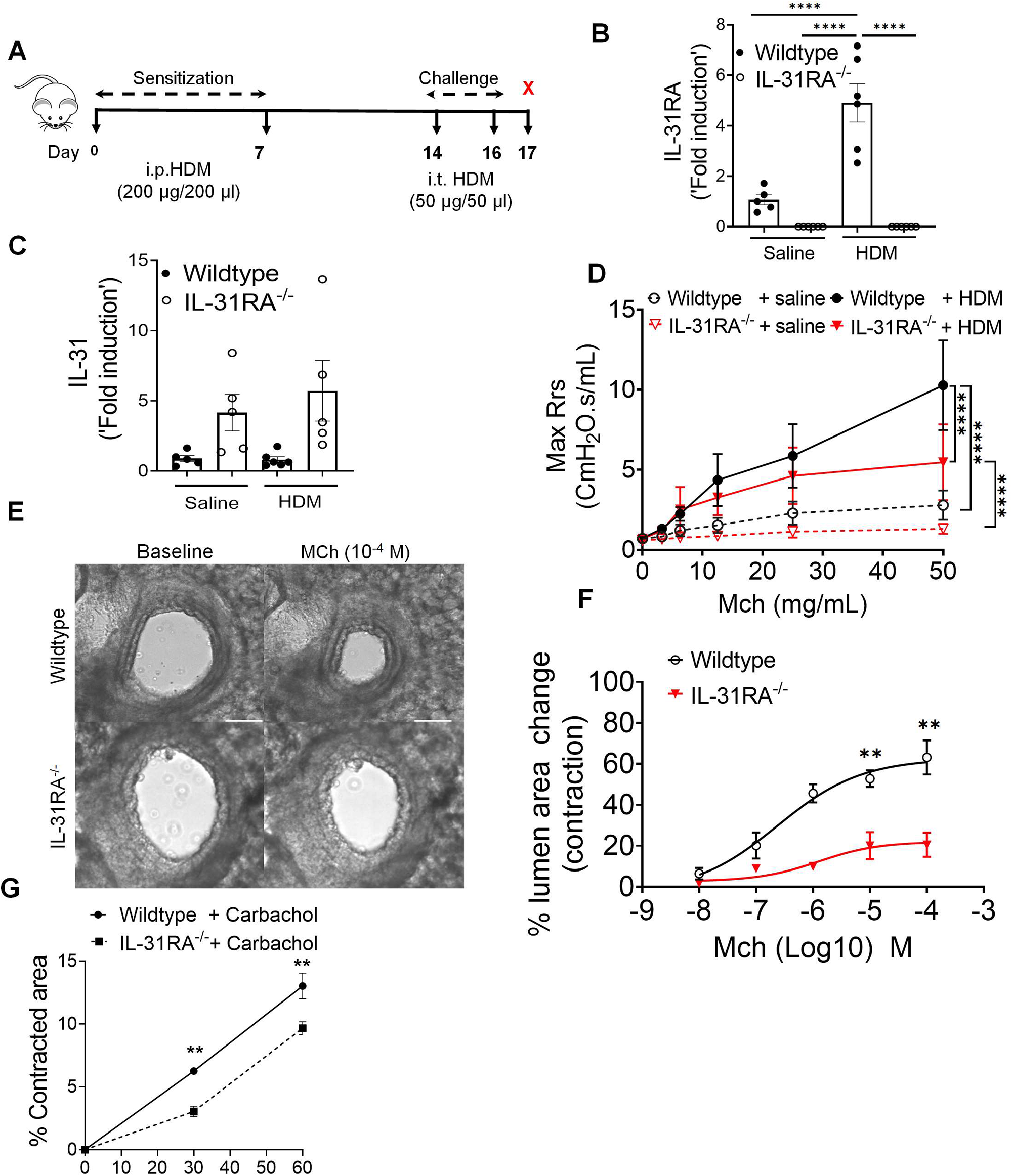
Loss of IL-31RA attenuates house dust mite (HDM)-induced airway hyperresponsiveness. **(A)** Schemata of HDM-induced allergic asthma model. **(B and C)** Quantification of IL-31RA and IL-31 transcripts in the lungs of wild-type and IL-31RA^-/-^ mice treated with HDM or saline. Data shown as mean ± SEM, n = 5-6/group, one-way ANOVA test, * p < 0.05. **(D)** Measurement of resistance with increasing doses of methacholine (MCh) in wild-type and IL-31RA^-/-^ mice treated with saline or HDM using FlexiVent. Data are shown as mean ± SEM, n = 7-8/group. A two-way ANOVA test, * p < 0.05. **(E)** Representative images of precision cut lung sections (PCLS) from the lungs of wild-type and IL-31RA^-/-^ mice treated with MCh (10^-4^ M) or prior to the treatment (baseline), scale bar 150 µm. **(F)** The percent of contraction of airways with increasing doses of MCh compared to the baseline area of airways between wild-type and IL-31RA^-/-^ mice. A two-way ANOVA test; n = 3-8 per group. * p < 0.05. **(G)** The percent contraction of collagen gels embedded with airway smooth muscle cells from wild-type and IL-31RA^-/-^ mice in culture media and treated with carbachol (10 µM) for 60 minutes. N = 3/group. Data shown as means mean ± SEM, two-tailed Student’s t-test, *p < 0.05

### Airway inflammation and goblet cell hyperplasia are IL-31RA independent

Mice primed and challenged with HDM develop strong Th2 responses, peribronchial inflammation, and goblet cell hyperplasia (*41, 42*). To determine whether IL-31RA was critical for the development of airway inflammation, we stained the lung sections of wild-type and IL-31RA^-/-^ mice challenged with HDM or saline with hematoxylin and eosin (H&E) (Fig 2A). As expected, the HDM challenge induced robust airway inflammation with immune cell infiltration in wild-type mice, as confirmed by quantitative measurement of the total bronchoalveolar lavage (BAL) cells (Fig 2B & 2C). Notably, the IL-31RA^-/-^ mice that were challenged with HDM developed airway inflammation similar to wild-type mice, indicating no effect of IL-31RA deficiency on allergic inflammatory response in airways. We further investigated whether the loss of IL-31RA had any effect on the infiltration of specific immune cell types into the airspaces by counting the immune cells in BAL. Consistent with the above findings, we observed no significant differences in the number of eosinophils, macrophages, lymphocytes, and neutrophils between wild-type and IL-31RA^-/-^ mice challenged with either saline or HDM (Fig 2C).

**Figure 2.**
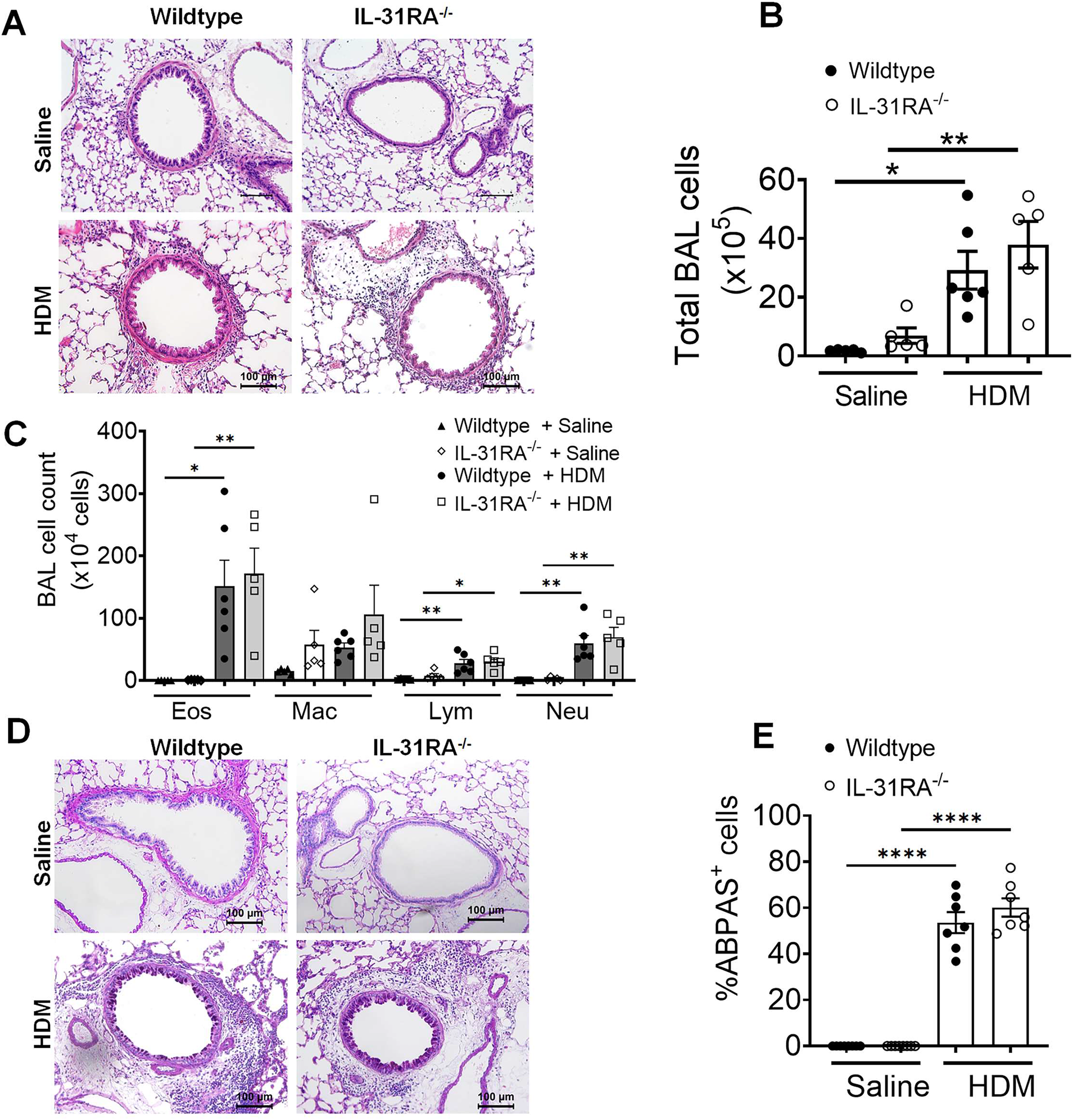
Loss of IL31RA has no effect on house dust mite (HDM)-induced airway inflammation and goblet cell hyperplasia. **(A)** Representative images of hematoxylin and eosin -stained lung sections from wild-type and IL-31RA^-/-^ mice treated with saline or HDM. Images were taken at 20x magnification, scale bar 100 µm. **(B and C)** Total bronchoalveolar lavage (BAL) cell number and the differential cell count of BAL cells of wild-type and IL-31RA^-/-^ mice treated with saline or HDM. Eos, eosinophils; Mac, macrophages; Lym, lymphocytes; and Neu, neutrophils. **(D)** Representative images of alcian blue periodic acid shiff(ABPAS) staining of lung sections from wild-type and IL-31RA^-/-^ mice treated with saline or HDM. Images were taken at 20x magnification, scale bar 100 µm. **(E)** The percent of ABPAS-positive cells normalized to total cell in the airways of wild-type and IL-31RA^-/-^ mice treated with saline or HDM. Data are shown as means ± SEM. One-way ANOVA was used, with n = 7-8/group. * p < 0.05.

Accumulation of mucus-secreting goblet cells or goblet cell hyperplasia is another important hallmark of allergic asthma observed in both human and animal models of allergic asthma (*43–45*). To determine whether there were differences in goblet cell hyperplasia, we performed alcian blue periodic acid-Schiff (ABPAS) staining to detect mucus-producing goblet cells in lung sections. Both wild-type and IL-31RA^-/-^ mice displayed increased goblet cell hyperplasia when challenged with HDM compared to unchallenged naïve animals (Fig 2D and 2E). These findings suggest that IL-31RA deficiency contributes to the development of AHR but has no significant effect on airway inflammation and mucus production in HDM-induced allergic asthma.

IL-31RA functions as a negative regulator of Th2 responses in the lungs (*46*) However, the role of IL-31RA in the development of Th2 responses during allergic asthma remains unclear. To determine whether there are alterations in the expression of asthma-associated genes in the absence of IL-31RA, we measured the transcript levels of Th2 cytokine, inflammatory cytokine, Th2 response, and goblet cell hyperplasia-associated genes. As expected, we observed a significant increase in Th2 cytokine gene expression, including IL-4, IL-5, and IL-13, in wild-type mice challenged with HDM compared to that with saline (Fig 3A). However, HDM-induced increase in Th2 cytokine expression was similar in the lungs of wild-type and IL-31RA^-/-^ mice, suggesting that the absence of IL-31RA had no effect on HDM-induced Th2 cytokine expression. Similarly, the loss of IL-31RA had no effect on the expression of chemokine and cytokine genes (CCL11, CCL24, and IL-10) and Th2 response-associated genes, including ARG1, CHI3L3, and FIZZ1 (Fig 3B & 3C). Consistent with ABPAS staining, we observed no effect of IL-31RA deficiency on HDM-induced expression of GOB5 and MUC5AC (Fig 3D). IL-6 and oncostatin M (OSM), together with IL-31, are members of the IL-6 family of cytokines known to regulate AHR in allergic asthma (*47–49*). Thus, we explored the levels of OSM and IL-6 in the lungs of wild-type and IL-31RA^-/-^ mice (Fig S1). We observed no significant differences in the expression of IL-6 and OSM between wild-type and IL-31RA^-/-^ mice treated with either saline or HDM, suggesting that the absence of IL-31RA did not affect the expression of OSM and IL-6.

**Figure 3.**
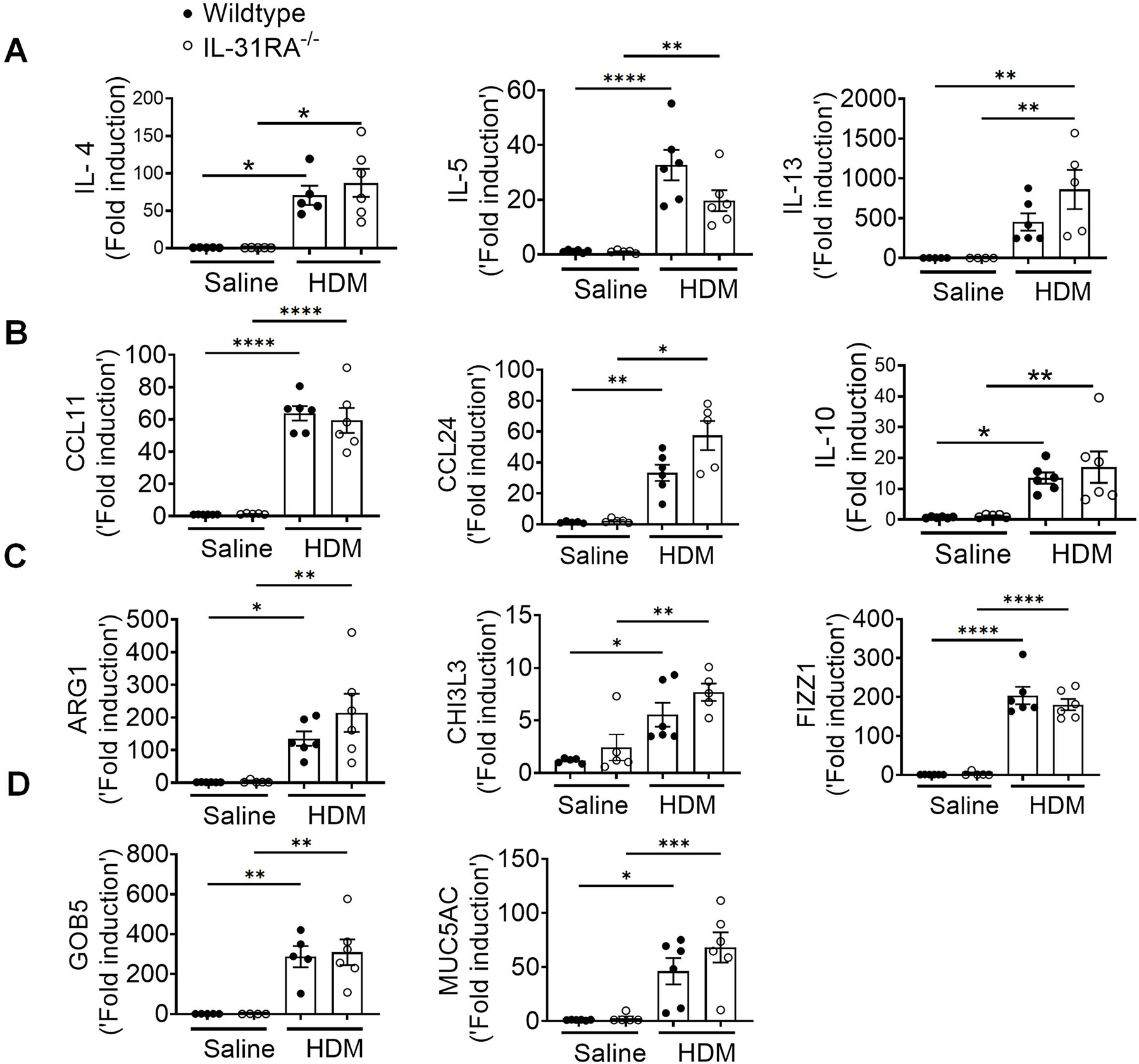
Loss of IL-31RA has no effect on Th2 responses and goblet cell hyperplasia during house dust mite (HDM)-induced allergic asthma. **(A)** Quantification of Th2 cytokine transcripts including IL-4, IL5, and IL-13 in the total lungs of wild-type and IL-31RA^-/-^ mice treated with saline or HDM using RT-PCR. **(B)** Quantification of chemokines and inflammatory cytokines including CCL11, CCL24, and IL-10 in the total lungs of wild-type and IL-31RA^-/-^ mice treated with saline or HDM using RT-PCR. **(C)** Quantification of Th2 response-associated gene transcripts including CHI3L3, ARG1, and FIZZ1 in the total lungs of wild-type and IL-31RA^-/-^ mice treated with saline or HDM using RT-PCR. **(D)** Quantification of GOB5 and MUC5AC gene transcripts in the total lungs of wild-type and IL-31RA^-/-^ mice treated with saline or HDM using RT-PCR. Data are shown as mean ± SEM and representative of two independent experiments with n=5-6/group. One-way ANOVA was used. * p < 0.05.

### IL-31RA is essential to induce AHR during SEA-induced allergic asthma

To further establish the role of IL-31RA in allergic asthma, we used an alternative mouse model of SEA-induced allergic asthma. Similar to HDM, extract of *Schistosoma mansoni* is a potent inducer of AHR, airway inflammation, and Th2 immune responses without the use of an adjuvant (*28, 50, 51*). As shown in the schematic, wild-type and IL-31RA^-/-^ mice were sensitized twice and consecutively challenged with SEA to induce allergic asthma (Fig 4A). The absence of IL-31RA gene expression was confirmed by measuring transcripts in the lungs of mice challenged with SEA or saline, and no IL-31RA expression was detected in IL31RA^-/-^ mice (Fig 4B). Consistent with the findings using the HDM model, sensitization and challenge with SEA resulted in a significant increase in the gene expression of IL-31RA but not IL-31 in the wild-type mice (Fig 4B & 4C).

**Figure 4.**
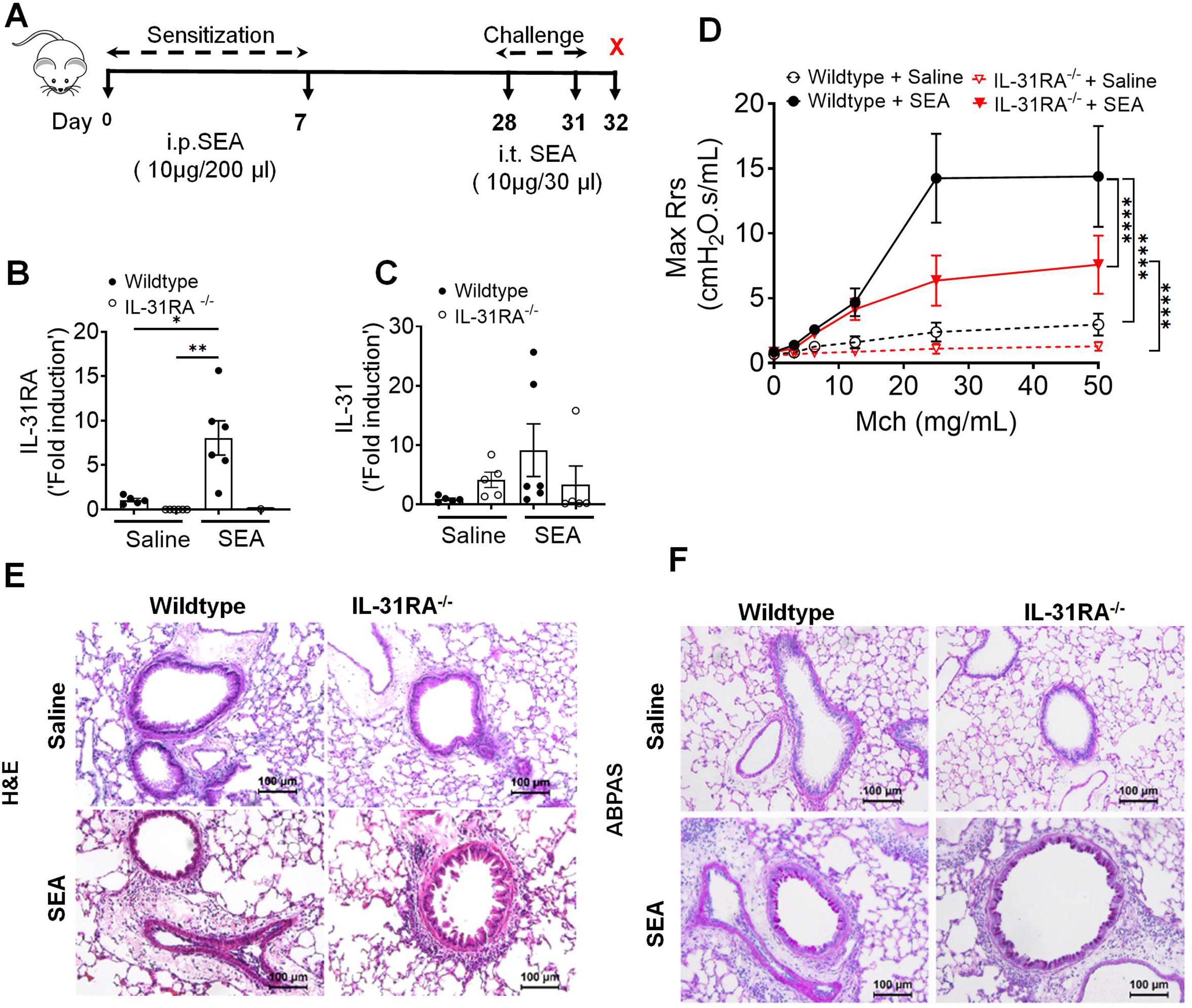
Loss of IL31RA attenuates airway hyperresponsiveness (AHR) but not inflammation and goblet cell hyperplasia during *Schistosoma mansoni* soluble egg antigen (SEA)-induced allergic asthma. **(A)** Schemata of SEA-induced allergic asthma model. **(B and C)** Quantification of IL-31RA and IL-31 transcripts in the lungs of wild-type and IL-31RA^-/-^ mice treated with house dust mite (HDM) or saline. Data shown as mean ± SEM, n = 5-6/group, one-way ANOVA test, *p < 0.05. **(D)** Measurement of resistance with increasing doses of methacholine (MCh) in wild-type and IL-31RA^-/-^ mice treated with saline or HDM using FlexiVent. Data are shown as mean ± SEM, n = 5-6/group. The above data is representative of two independent experiments with similar results. A two-way ANOVA test for multiple comparisons was used. * p<0.01. **(E)** Representative images of hematoxylin and eosin-stained lung sections from wild-type and IL-31RA^-/-^ mice treated with saline or SEA. Images were taken at 20x magnification, scale bar 100 µm. **(F)** Representative images of alcian blue periodic acid shiff staining of lung sections from wild-type and IL-31RA^-/-^ mice treated with saline or SEA. Images were taken at 20x magnification, scale bar 100 µm.

To assess whether the loss of IL-31RA attenuates SEA-induced AHR, we measured MCh-induced AHR in both wild-type and IL-31RA^-/-^ mice sensitized and challenged with either saline or SEA (Fig 4D). Loss of IL-31RA was sufficient to attenuate MCh-induced AHR in SEA-challenged mice (Fig 4D). To evaluate the changes in airway inflammation, lung sections of wild-type and IL31RA^-/-^ mice challenged with SEA or saline were stained with H&E. As observed in the HDM model, SEA challenge induced robust airway inflammation with immune cell infiltration in both wild-type and IL-31RA^-/-^ mice (Fig 4E). Similarly, the airways of IL-31RA^-/-^ mice that were challenged with SEA showed an accumulation of mucus-producing goblet cells similar to wild-type mice, as seen in ABPAS-stained lung sections (Fig 4F).

To determine whether IL-31RA-induced AHR is regulated by asthma-associated gene networks, we measured the expression of Th1 and Th2 cytokines. The expression of IFN-γ IL-4, and IL-5 was elevated in both wild-type and IL-31RA^-/-^ mice following sensitization and challenge with SEA (Fig 5A). Similarly, the transcripts of genes associated with inflammation (CCL11, CCL24, and IL-10) and Th2 responses (ARG1, CHI3L3, and FIZZ1) remained elevated in IL-31RA^-/-^ mice, and no differences were observed when compared to wild-type mice treated with SEA (Fig 5B & 5C). The gene transcripts associated with goblet cell hyperplasia were elevated in both wild-type and IL-31RA^-/-^ mice treated with SEA (Fig 5D). In summary, in the two alternative mouse models of allergic asthma, the absence of IL-31RA altered AHR but did not appear to affect airway inflammation, Th2 responses, or goblet cell hyperplasia.

**Figure 5.**
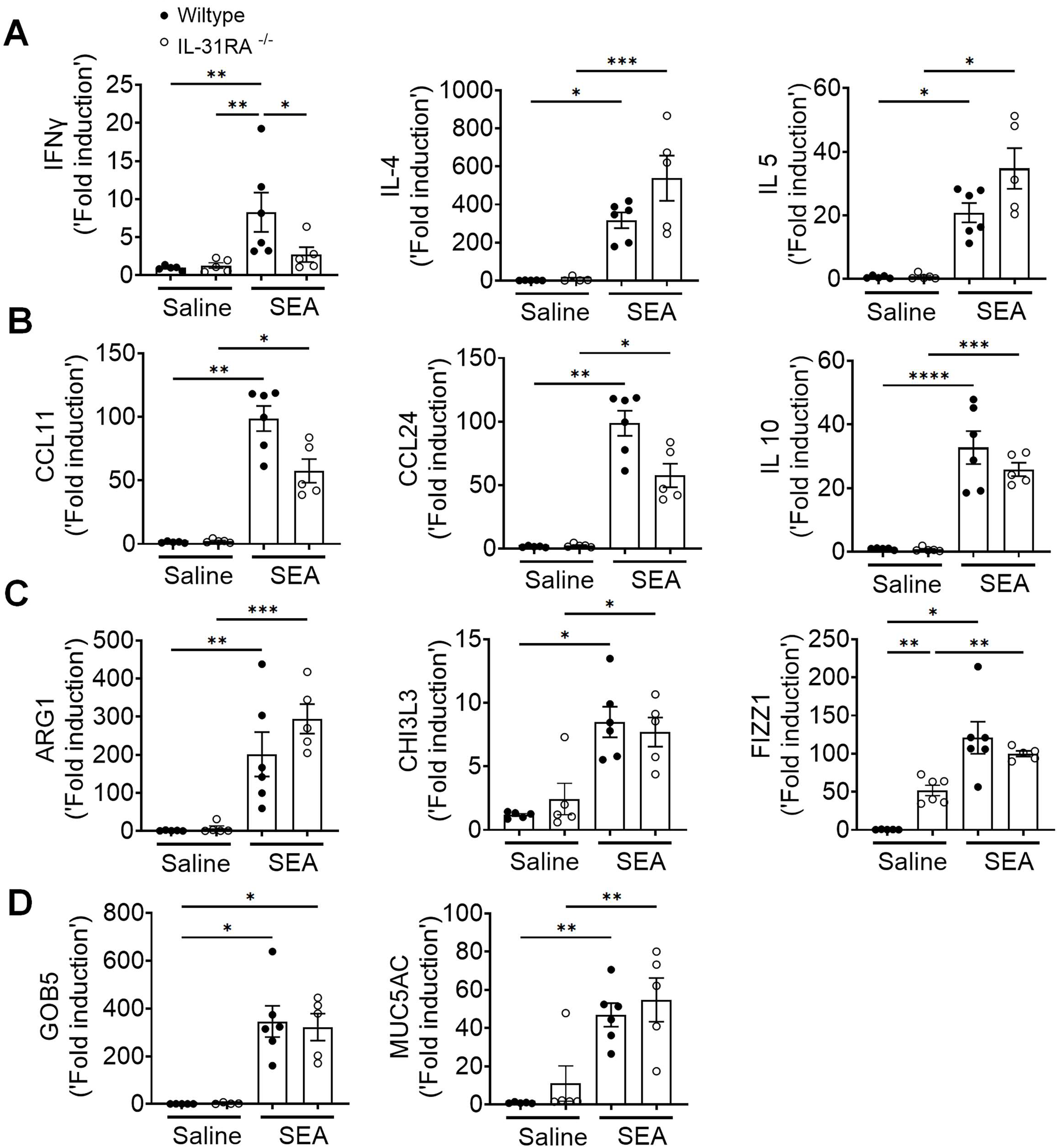
Loss of IL31RA has no effect on Th2 responses and goblet cell hyperplasia during *Schistosoma mansoni* soluble egg antigen (SEA)-induced allergic asthma. **(A)** Quantification of IFN-γ, IL-4, IL-5, and IL-13 in the total lungs of wild-type and IL-31RA^-/-^ mice treated with saline or SEA using RT-PCR. **(B)** Quantification of CCL11, CCL24, and IL-10 in the total lungs of wild-type and IL-31RA^-/-^ mice treated with saline or SEA using RT-PCR. **(C)** Quantification of Th2 response-associated genes including CHI3L3, ARG1, and FIZZ1 in the total lungs of wild-type and IL-31RA^-/-^ mice treated with saline or SEA using RT-PCR. **(D)** Quantification of GOB5 and MUC5AC gene transcripts in the total lungs of wild-type and IL-31RA^-/-^ mice treated with saline or SEA using RT-PCR. Data are shown as means ± SEM and representative of two independent experiments with n=5-6/group. One-way ANOVA was used. * p < 0.05.

### IL-31 is dispensable to induce AHR, inflammation, and Th2 responses

IL-31RA is a cognate binding receptor of IL-31 that recruits OSMRβ to form a high-affinity binding receptor complex to signal through the JAK/STAT pathway (*29, 30*). Previous studies have demonstrated that increased IL-31 signaling through IL-31RA can result in uncontrolled inflammation and tissue remodeling in multiple tissues, including the lungs and skin (*25, 26*). However, we observed no significant increase in IL-31 levels in the lungs during HDM- and SEA-induced allergic asthma. To assess the potential pathological effects of IL-31 in asthma, the lungs of wild-type mice were intratracheally treated with saline or IL-31, and changes in AHR and inflammation were assessed (Fig 6A). Notably, we observed no significant changes in AHR between saline and IL-31 treated wild-type mice (Fig 6B). Similarly, we observed no significant changes in tissue inflammation as assessed by H&E-stained lung sections of saline and IL-31 treated wild-type mice (Fig 6C). To confirm the signaling effects of IL-31, we measured the expression of the IL-31-driven gene suppressor of cytokine signaling 3 (*SOCS3*) in the lungs of mice treated with saline and IL-31. Previously published studies have shown that IL-31 upregulates *SOCS3* (52); similarly, the expression of *SOCS3* was significantly upregulated in lungs of IL-31-treated mice compared to the saline-treated mice (Fig 6D). To determine the effects of IL-31 on the expression of asthma-associated genes, we quantified the expression of genes associated with inflammation (*IFN-γ, TNF-α, IL-6, and IL-17*) and Th2 responses (*IL-4, IL-13, ARG1, MUC4, and MUC5AC*). Notably, we observed no significant changes in the expression of genes associated with either inflammation or Th2 responses (Fig 6E and 6F).

**Figure 6.**
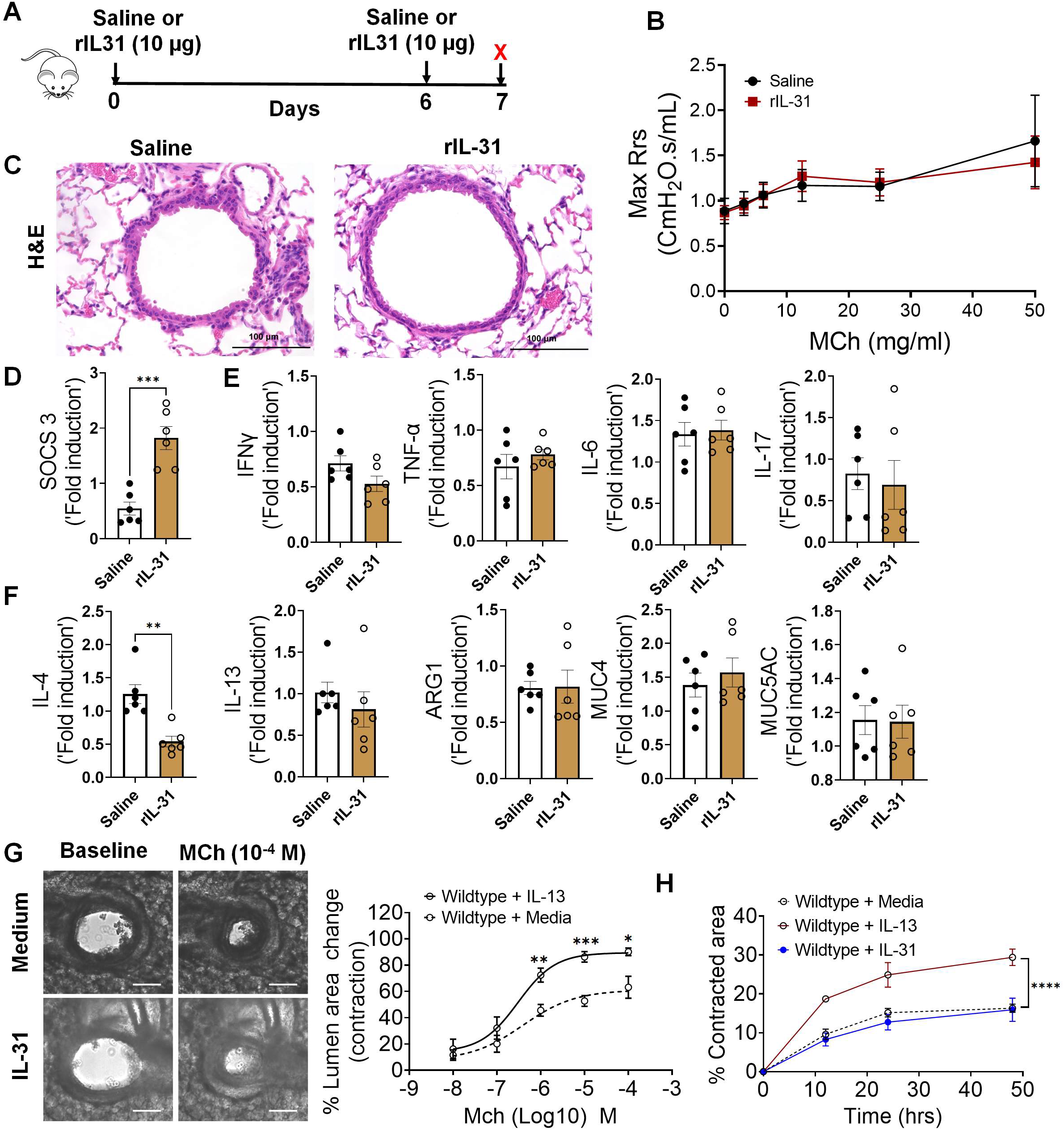
IL-31 is dispensable for the induction of airway hyperresponsiveness (AHR), inflammation, and Th2 responses. **(A)** Schemata showing intratracheal administration of IL-31 or saline in wild-type mice. **(B)** Measurement of resistance with increasing doses of methacholine (MCh) in wild-type mice treated with saline or IL-31 using FlexiVent. Data are shown as mean ± SEM, n = 6/group. The data is representative of two independent experiments with no statistical significance between groups. **(C)** Representative images of hematoxylin and eosin-stained lung sections from wild-type mice treated with IL-31 or saline. Images were captured at 20x magnification, scale bar 100 µm. (D) Quantification of IL-31-induced *SOCS3* gene expression in the total lungs of wild-type mice treated with saline or IL-31. Unpaired two-tailed Student’s t-test was used. ***P < 0.001. (E and F) Quantification of inflammation-associated gene transcripts *IFN-γ TNFα IL-6,* and *IL-17* and *Th2*-associated gene transcripts including *IL-4, IL-13, ARG1, MUC4*, and *MUC5AC* in the total lungs of wild-type mice treated with saline or IL-31. Unpaired two-tailed Student’s t-test was used and no significance found between groups, n=6/group. (G) Representative images of precision cut lung sections (PCLS) from wild-type mice treated with or without IL-31 (500 ng/ml) for 24 h. Airway contractility was measured in response to MCh (10^-4^ M) compared to baseline diameter. The percent of airway lumen area contraction with increasing doses of MCh was calculated for saline and IL-31-treated PCLS from wild-type mice. Two-way ANOVA test, n=5-8/group. (H) The percent contraction of collagen gels embedded with airway smooth muscle cells from wild-type mice that were treated with media, IL-13 (50 ng/ml) or IL-31 (500 ng/ml). The percent contraction was measured at different time points compared to baseline. Two-way ANOVA was used, n=4/group.

Furthermore, we evaluated airway contractility induced by IL-31 using PCLS. As demonstrated in previous studies, we observed a significant increase in the contractility of airways of wild-type mice treated with IL-13 compared to saline-treated mice in a dose-dependent manner with MCh (Fig S2). Conversely, we observed no significant effect of IL-31 on the contractility of wild-type airways exposed to MCh in comparison to media (Fig 6G). Furthermore, we observed no significant changes in the kinetics of contraction of collagen gels embedded with ASMC, which were treated with either media alone or IL-31. However, we observed a significant increase in the contraction of collagen gels embedded with ASMC and treated with IL-13 compared to that with media alone (Fig 6H). Thus, in contrast to IL-31RA, the results suggest that IL-31 is not involved in modifying AHR, inflammation, and Th2 responses in the lungs; the reduced AHR observed in the absence of IL-31RA could be due to other alternative mechanisms that need to be identified.

### Th2 cytokines upregulate IL-31RA to induce AHR with no effect on inflammation

Previous studies from our lab and others have shown that both IL-4 and IL-13 induce IL-31RA expression in macrophages and lungs via the type II IL-4 receptor and STAT6 signaling (*38*). To determine whether IL-4 or IL-13 can induce IL-31RA expression in ASMC, we treated ASMC with increasing doses of IL-4 or IL-13 and quantified the expression of IL31RA. Consistent with our earlier findings in macrophages, the expression of IL-31RA significantly increased in ASMC treated with IL-4 or IL-13 compared to that with media (Fig 7A & 7 B). IL-13 is a potent inducer of AHR, inflammation, and goblet cell hyperplasia in mice (*18*). To determine whether IL-13-induced AHR is IL-31RA dependent, we treated both wild-type and IL-31RA^-/-^ mice with IL-13 and assessed AHR and other pathological and molecular changes that are relevant to asthma. Notably, the loss of IL-31RA was sufficient to attenuate IL-13-induced AHR compared to wild-type mice exposed to increasing doses of MCh (Fig 7C). Similarly, the contraction of collagen gels embedded with ASMC from IL-31RA^-/-^ mice significantly reduced compared to wild-type mice with IL-13 treatment (Fig 7D). In contrast, H&E staining of lung sections suggested that the loss of IL-31RA had no effect on IL-13-driven airway inflammation in wild-type and IL31RA^-/-^ mice (Fig 7E). Quantitative assessment of inflammatory chemokines and cytokines suggested that the loss of IL-31RA had no effect on the expression of CCL11, CCL24, and IL-17 (Fig 7F). Similarly, we observed no defects in the expression of Th2 cytokines (IL-4 and IL-5) or genes associated with Th2 responses (ARG1, CHI3L3, and FIZZ1) in IL-31RA^-/-^ mice compared to wild-type mice treated with IL-13 (Fig 7G and 7H).

**Figure 7.**
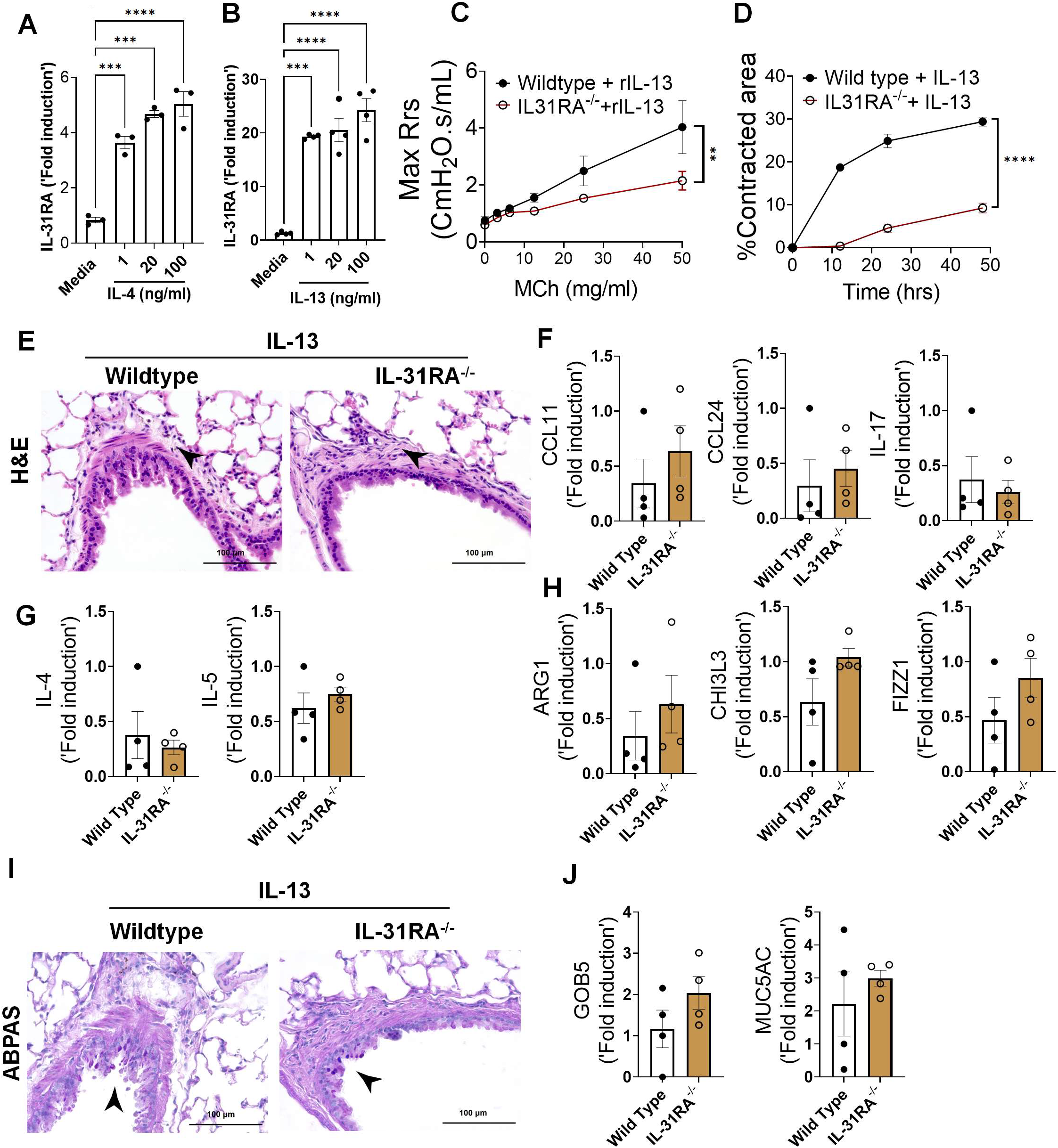
Th2 cytokines upregulate IL-31RA to induce airway hyperresponsiveness (AHR) with no effect on inflammation and goblet cell hyperplasia. **(A and B)** Quantification of IL-31RA transcripts in mouse airway smooth muscle cells (ASMC) treated with increasing doses of IL-4 and IL-13 for 16 h. One-way ANOVA test was used, n = 3/group; * p<0.05. **(C)** Wild-type and IL-31RA^-/-^ mice were treated intratracheally with IL-13 on days D0 and D6 and resistance in the lungs was measured with increasing doses of methacholine (MCh) using FexiVent. Data are shown as mean ± SEM, n=4/group. Two-way ANOVA test was used, *p<0.05. **(D)** ASMC isolated from wild-type and IL-31RA^-/-^ mice were seeded into collagen gels and treated with IL-13 to measure the contraction of collagen gels after 48 hours. Two-tailed Student’s t-test was used, n=3/group; * p<0.05. **(E)** Representative images of hematoxylin and eosin -stained lung sections from wild-type and IL-31RA^-/-^ mice treated with IL-13. Images were captured at 20x magnification, scale bar 100 µm. **(F, G and H)** Quantification of inflammatory chemokines (*CCL11, CCL24 and IL-17,* Th2 cytokines (*IL-4* and *IL-5*), and Th2 response-associated genes including *ARG1, CHl3L3,* and *FIZZ1* in the total lungs of wild-type and IL-31RA^-/-^ mice treated with IL-13. Data are shown as mean ± SEM, n=4/group, Two-tailed Student’s t-test was used and no statistical significance observed between groups. **(I)** Representative images of alcian blue periodic acid shiff stained lung sections from wild-type and IL-31RA^-/-^ mice treated with IL-13. Images were captured at 20x magnification, scale bar 100 µm. **(J)** Quantification of goblet cell hyperplasia associated genes including *GOB5* and *MUC5AC* transcript levels in the total lungs of wild-type and IL-31RA^-/-^ mice treated with IL-13. Data are shown as means ± SEM, n=4/group, Two-tailed Student’s t-test was used and no statistical significance observed between groups.

To evaluate whether IL-13-induced goblet cell hyperplasia was altered in the absence of IL-31RA, we assessed the accumulation of goblet cells in the airways and quantified the transcripts associated with goblet cell hyperplasia. As shown in figure 7I, goblet cell accumulation was similar between wild-type and IL31RA^-/-^ mice treated with IL-13. In support of this, the expression of *GOB5* and *MUC5AC* remained similar between IL-13 treated wild-type and IL31RA^-/-^ mice (Fig 7J). Thus, IL-31RA expression is essential to mediate IL-13-induced AHR but dispensable for inflammation and goblet cell hyperplasia.

### IFN-γ upregulates IL-31RA to induce AHR with no effects on inflammation

IFN-γ is a key Th1 cytokine implicated in elevated AHR in patients with severe asthma or mixed Th1/Th2 cytokine phenotypes (*12, 53*). To determine whether IFN-γ induces the expression of IL-31RA in ASMC, we isolated ASMC from the trachea of wild-type mice and treated them with increasing doses of IFN-γ The expression of IL-31RA is significantly increased by IFN-γ in a dose dependent manner (Fig 8A). A recent study suggested that IL-31 can also upregulate the expression of IL-31RA in human sensory neurons (*54*). However, we observed no significant effect of IL-31 on the expression of IL-31RA in ASMC (Fig 8B). To determine the effects of IFN-γ on asthma phenotypes, we treated wild-type mice with saline or IFN-γ and measured the changes in AHR, inflammation, and goblet cell hyperplasia. We observed a significant increase in AHR in wild-type mice treated with IFN-γ compared to that with saline (Fig 8C). In addition, we observed a significant increase in peri-bronchial inflammation in wild-type mice treated with IFN-γ compared to that with saline (Fig 8D). To establish the quantitative gene expression changes induced by IFN-γ, we measured IRF1, IRF7, and STAT1 and observed a significant increase in the expression of IFN-γ -specific genes in the lungs (Fig S3).

**Figure 8.**
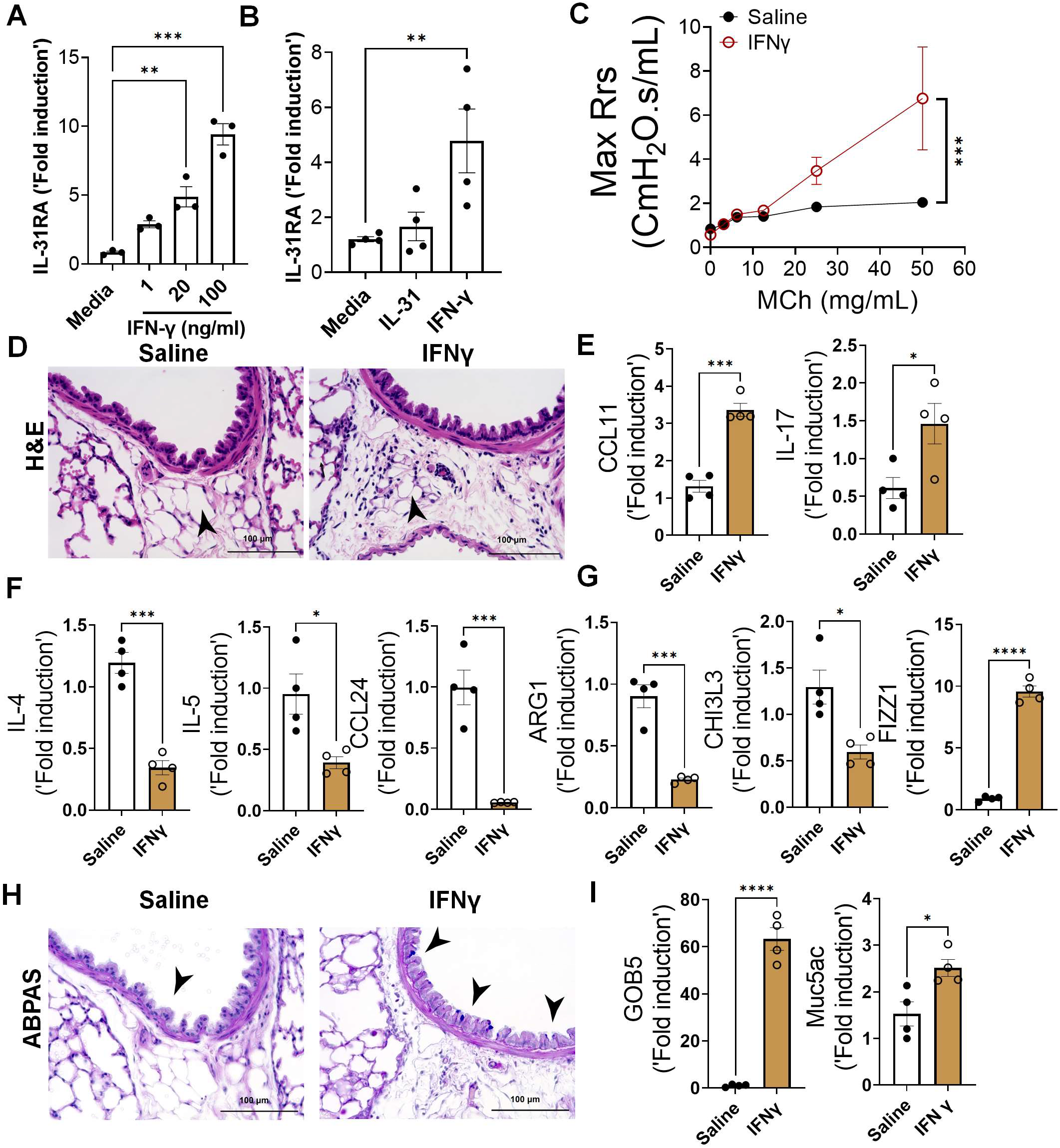
IFN-γ is a positive regulator of IL31RA expression, along with airway hyperresponsiveness (AHR), inflammation, and goblet cell hyperplasia. **(A)** Quantification of IL-31RA transcript levels in mouse airway smooth muscle cells (ASMC) treated with increasing doses of IFN-γ for 16 h. Data are shown as the mean ± SEM (n = 3/group). One-way ANOVA was used; *P < 0.05. **(B)** Quantification of IL-31RA transcript levels in human ASMCs treated with IL-31 (500 ng/ml) or IFN-γ (50 ng/ml) for 16 h. Data are shown as the mean ± SEM, n = 3-4/group. One-way ANOVA was used; *P < 0.05. **(C)** Wild-type mice were intratracheally treated with IFN-γ (5 µg) on days 0 and 6, and resistance to increasing doses of methacholine was measured on day 7. Data are shown as mean ± SEM, n = 4/group; two-way ANOVA was used, * P < 0.05. **(D)** Representative images of hematoxylin and eosin -stained lung sections from wild-type mice treated with saline or IFN-γ. Images were captured at 20x magnification, with a scale bar of 100 µm. **(E, F, and G)** Quantification of inflammatory chemokines (*CCL11* and *IL-17*), Th2 cytokines (*IL-4, IL-5*, and *CCL24*), and Th2 response-associated genes, including *ARG1*, *CHl3L3*, and *FIZZ1*, in the lungs of wild-type mice treated with saline or IFN-γ. Data are shown as the mean ± SEM, n = 6/group. Two-tailed Student’s t-test was used; * P < 0.05. **(H)** Representative images of alcian blue periodic acid shiff -stained lung sections from wild-type mice treated with saline or IFN-γ. Images were captured at 20x magnification, scale bar = 100 µm. **(I)** Quantification of mucus-associated gene transcripts, including *GOB5* and *MUC5AC*, in the lungs of wild-type mice treated with saline or IFN-γ. Data are shown as the mean ± SEM (n = 6/group). Two-tailed Student’s t-test was used; * P < 0.05.

To determine the effects of IFN-γ on asthma phenotypes, we measured the expression of genes associated with inflammation and Th2 responses. In support of the substantial inflammation observed after IFN-γ treatment, we observed a significant increase in CCL11 and IL-17 expression (Fig 8E). As anticipated, IFN-γ treatment resulted in negative regulation of Th2 cytokines and Th2 response-associated gene expression, including IL-4, IL-5, CCL24, ARG1, and CHI3L3, but had no effect on the expression of FIZZ1, which was significantly increased compared to the saline group (Fig 8F and G). Similar to IL-13, IFN-γ was able to induce the accumulation of goblet cells in the airways with increased expression of Gob5 and MUC5AC (Fig 8H and 8I). Despite the reduced expression of several Th2-associated asthma genes, IFN-γ treatment induced AHR, inflammation, and goblet cell hyperplasia.

To determine whether the expression of IL-31RA was critical for IFN-γ -induced AHR, inflammation, and goblet cell hyperplasia, we intratracheally treated both wild-type and IL31RA^-/-^ mice with IFN-γ. Notably, IFN-γ -induced AHR was significantly attenuated in IL31RA^-/-^ mice compared to that in wild-type mice (Fig 9A and Fig S4). However, as noted with IL-13, increases in peribronchial inflammation and the expression of inflammatory cytokine genes, such as CCL11, CCL24, and IL-17, were similar between wild-type and IL-31RA^-/-^ mice treated with IFN-γ Fig 9B and 9C). Furthermore, we observed no significant differences in the expression of Th2 cytokine genes (IL-4 and IL-5) or the expression of genes associated with Th2 responses, including ARG1, CHI3L3, and FIZZ1 (Fig 9D and 9E).

**Figure 9.**
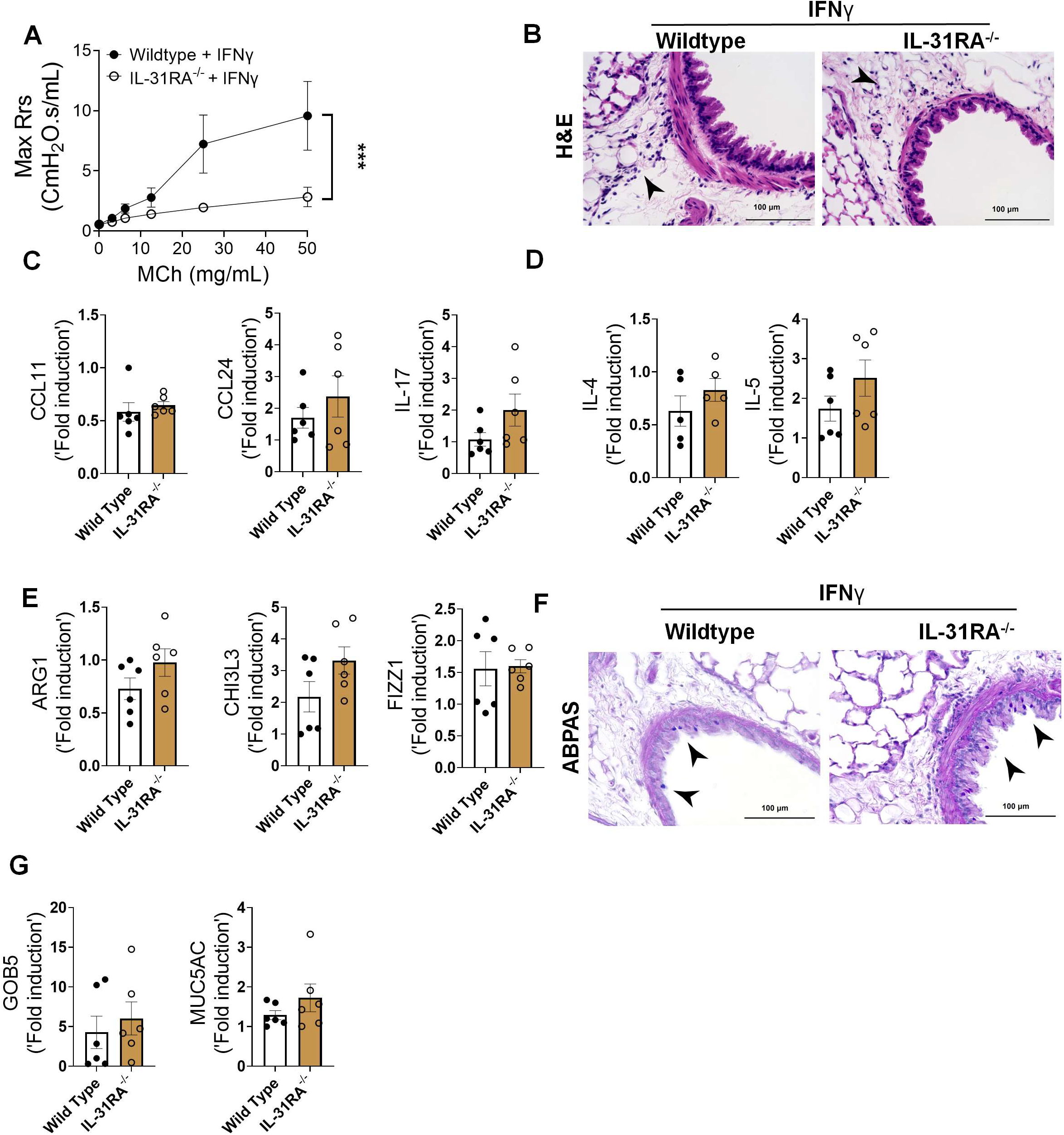
Loss of IL-31RA is sufficient to attenuate IFN-γ -driven airway hyperresponsiveness (AHR) but no effect on inflammation and goblet cell hyperplasia. **(A)** Wild-type and IL-31RA^-/-^ mice were treated intratracheally with IFN-γ on days D0 and D6 and resistance in the lungs was measured with increasing doses of methacholine (MCh) using FexiVent. Data are shown as mean ± SEM, n = 6/group. Two-way ANOVA was used, *p<0.05. **(B)** Representative images of hematoxylin and eosin -stained lung sections from wild-type and IL-31RA^-/-^ mice treated with IFN-γ were captured at 20x magnification, scale bar 100 µm, n = 6/group. **(C, D and E)** Quantification of inflammatory cytokines (*CCL11, CCL24* and *IL-17),* Th2 cytokines (*IL-4* and *IL-5*), and Th2 response-associated genes including *ARG1, CHl3L3,* and *FIZZ1* in the lungs of wild-type and IL-31RA^-/-^ mice treated with IFN-γ. Data are shown as mean ± SEM, n = 6/group. Two-tailed Student’s t-test was used and no statistical signification observed between groups. **(F)** Representative images of alcian blue periodic acid shiff-stained lung sections from wild-type and IL-31RA^-/-^ mice treated with IFN-γ. Images were captured at 20x magnification, scale bar 100µm, n = 6/group. (G) Quantification of mucus-associated genes including *GOB5* and *MUC5AC* transcript levels in the lungs of wild-type and IL-31RA^-/-^ mice treated with IFN-γ. Data are shown as means ± SEM, n = 6/group. Two-tailed Student’s t-test was used, and no statistical signification observed between groups.

To evaluate the effects of IL-31RA deficiency on IFN-γ -induced goblet cell hyperplasia, we performed ABPAS staining of lung sections from wild-type and IL-31RA^-/-^ mice and quantified GOB5 and MUC5AC expression. Analysis of ABPAS-stained lung sections suggested no change in the number of goblet cells that accumulated in the airways of wild-type and IL31RA^-/-^ mice treated with IFN-γ Fig 9F). This finding was further substantiated by quantitative PCR data, which showed no quantitative differences in the expression of GOB5 and MUC5AC with the loss of IL-31RA (Fig 9G). Nevertheless, our in vivo studies demonstrated that despite the development of substantial inflammation and mucus hypersecretion, IL-31RA^-/-^ mice showed attenuated AHR in response to either IL-13 or IFN-γ. Overall, our new finding convincingly demonstrate that in allergic asthma, IL-31RA, which is induced by both Th1 and Th2, is critically required for the development of AHR but not inflammation and mucus secretion.

### IL-31RA augments CHRM3-driven calcium signaling in ASMC

Muscarinic acetylcholine receptors (CHRMs) are predominantly expressed by structural cells such as smooth muscle cells of airways and play a major role in triggering contraction of airways and AHR (*55, 56*). To evaluate mechanisms by which IL-31RA induces AHR, we measured the transcripts of five major receptor subtypes including CHRM1, CHRM2, CHRM3, CHRM4 and CHRM5 in the lungs of wild-type and IL-31RA^-/-^ mice. Quantification of the lung transcripts suggest no significant differences in the transcript levels of CHRMs in the lungs of IL-31RA^-/-^ mice compared to wild-type mice (Fig 10A). CHRM3 is a dominant receptor subtype expressed in ASMC and coupled to the G_q/11_ family of G proteins to induce calcium signaling and the contraction of ASMC in asthma(55, 57). Therefore, we evaluated the changes in the transcripts of CHRM3 by IL-4 and IL-31 in ASMC of wild-type and IL-31RA^-/-^ mice. Both IL-4 and IL-31 had little or no effect on the expression of CHRM3 either in the presence or absence of IL-31RA suggesting other mechanisms might be involved in IL-31RA-driven contraction of ASMC (Fig 10B and 10C). To determine post-transcriptional regulation of CHRM3, we measured the protein levels of CHRM3 in AMSC isolated from wild-type and IL-31RA^-/-^ mice. Notably, we observed a significant decrease in the protein levels of CHRM3 in ASMC isolated from IL-31RA^-/-^ mice compared to wild-type mice (Fig 10D). The decrease in CHRM3 protein with no changes in the transcript levels in the absence of IL-31RA may suggest a post-transcriptional stabilization of CHRM3 that may involve a physical interaction between IL-31RA and CHRM3 in ASMC. To identify the complex formation between IL-31RA and CHRM3, we used *in situ* proximity ligation method and measured the complex formation between IL-31RA and CHRM3 in HEK293 overexpressing them. We generated overexpression plasmids and our western blot analysis of HEK293 cell lysates show overexpression of IL-31RA and CHRM3 in HEK293 cells (Fig S5). Importantly, we observed bright fluorescent signals corresponding to the IL-31RA-CHRM3 complex formation in HEK293 cells overexpressing IL-31RA and CHRM3 compared to control cells (Fig 10E). By fluorescently labeling plasma membrane with cholera toxin, we demonstrated the colocalization of PLA puncta with the plasma membrane in HEK293 cells overexpressing IL-31RA and CHRM3 compared to control cells (Fig S6). These results demonstrated the specificity of the PLA methodology to detect IL-31RA-CHRM3 complex formation selectively in HEK293 cells overexpressing both IL-31RA and CHRM3 compared to control cells. Next, we assessed the gain-of-function effects of IL-31RA on CHRM3-driven calcium signaling by carbachol in HEK293 cells. Carbachol treatment in HEK293 cells transfected with control plasmids resulted in elevated intracellular calcium as these cells naturally express low levels of CHRM3 (Fig 10F). Notably, overexpression of IL31RA was sufficient to augment calcium release similar to the levels observed with CHRM3 overexpression. This increase in intracellular calcium release was further elevated in HEK293 cells that co-expressed both IL-31RA and CHRM3 which may suggest a cooperation between IL-31RA and CHRM3 to augment carbachol-induced intracellular calcium. However, calcium-dependent phosphorylation of myosin light chain (MLC) is a terminal event in the contraction of ASMC (*58*). Therefore, we measured carbachol-induced phosphorylation of MLC in HEK293 cells overexpressing IL-31RA compared to control HEK293 cells. Importantly, overexpression of IL-31RA alone was sufficient to augment the levels of carbachol-induced MLC phosphorylation that involved in smooth muscle contraction (Fig 10G). To further demonstrate the positive regulation of MLC phosphorylation by the IL-31RA-CHRM3 axis, we measured carbachol-induced phosphorylation of MLC in ASMC isolated from wild-type and IL-31RA^-/-^ mice. Consistent with reduced CHRM3 expression in IL-31RA deficient ASMC, we observed a significant decrease in carbachol-induced MLC phosphorylation in ASMC isolated from IL-31RA^-/-^ mice compared to wild-type mice (Fig 10H). These results suggest that IL-31RA functions as a positive regulator of CHRM3 and associated carbachol-induced calcium signaling to augment the contractility of ASMC in asthma.

**Figure 10.**
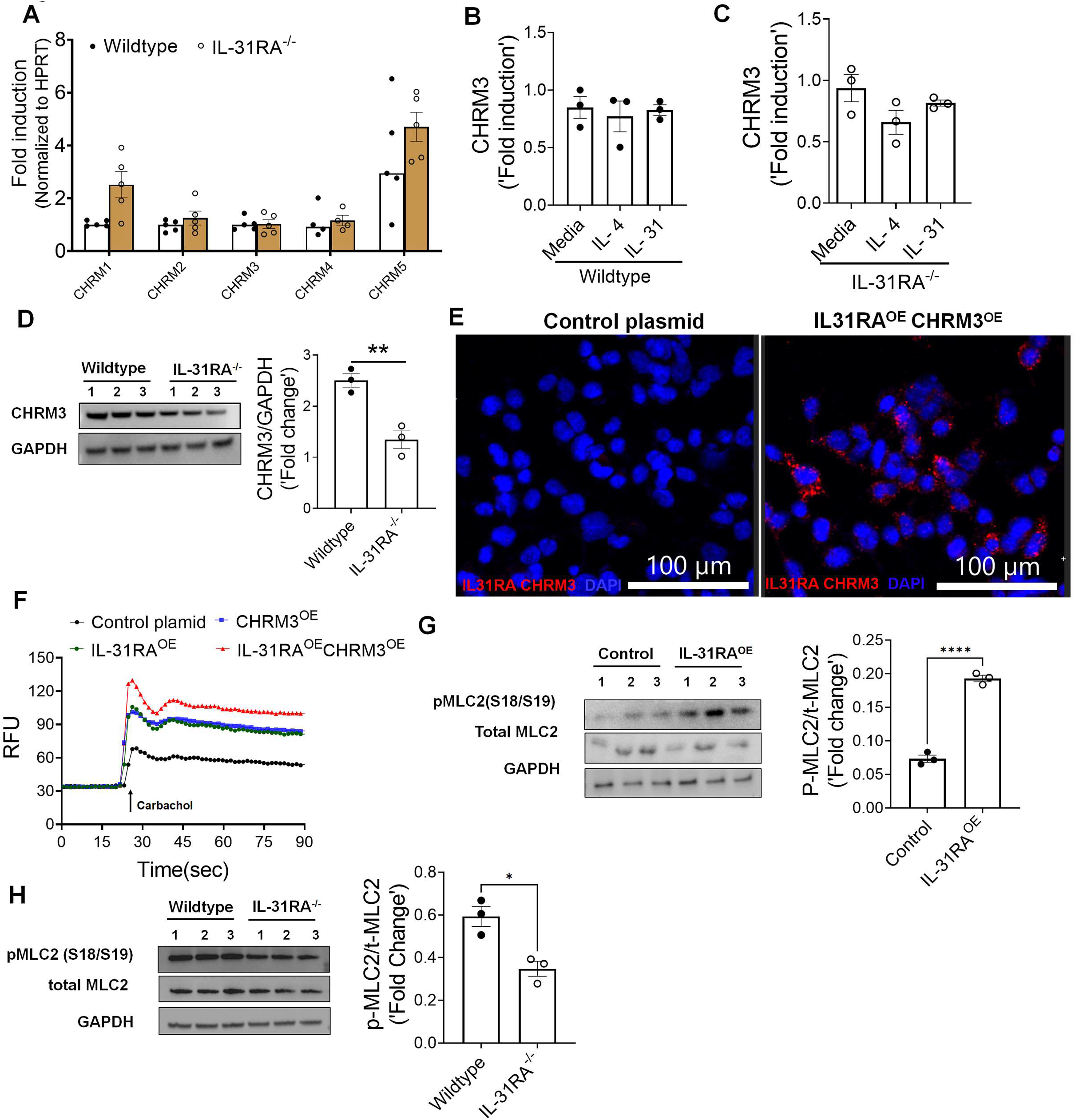
IL-31RA augments CHRM3-driven calcium signaling and MLC phosphorylation in ASMC. **(A)** Quantification of the transcripts of different isoforms of CHRMs in the lungs of wild-type and IL-31RA^-/-^ mice. **(B)** Quantification of the transcripts of CHRM3 in ASMC isolated from wildtype mice and treated with media, IL-4 (10 ng/ml) or IL-31 (500 ng/ml) for 16 h. **(C)** Quantification of the transcripts of CHRM3 in ASMC isolated from IL-31RA^-/-^ mice and treated with media, IL-4 (10 ng/ml) and IL-31 (500 ng/ml) for 16 h. **(D)** ASMC isolated from wild-type and IL-31RA^-/-^ mice were lysed and immunoblotted with antibodies against CHRM3 and GAPDH. CHRM3 protein levels were normalized to GAPDH and shown as fold induced using a bar graph (***P < 0.0005; n = 3, Student’s 2-tailed t test). **(E)** HEK293 cells were transiently co-transfected with overexpressing plasmids for CHRM3 and IL-31RA or empty control plasmids for 48 h. The IL-31RA-CHRM3 complex formation was visualized using hybridization probes labeled with Alexa 594 (Red). The nuclei were stained with DAPI (blue) and images were captured at 40X magnification. Scale bar, 100 µm. **(F)** Increases in intracellular calcium levels were measured in HEK 293 cells transiently transfected with overexpressing plasmids for CHRM3 and IL-31RA or empty control plasmids for 72 h and treated with carbachol (10µM). **(G)** HEK 293 cells were transiently co-transfected with overexpressing plasmids for CHRM3 and IL-31RA or empty control plasmids for 48 h and treated with carbachol (10µM) for 10 min. Cell lysates were immunoblotted with antibodies against phospho-MLC, total-MLC and GAPDH. Data are shown as means ± SEM, n = 3/group. **(H)** ASMC isolated from wild-type and IL-31RA^-/-^ mice were treated with carbachol (10µM) for 10 min and cell lysates were immunoblotted with antibodies against phospho-MLC, total-MLC and GAPDH. Data are shown as means ± SEM, n = 3/group

## DISCUSSION

In this study, we investigated the pathophysiological role of the IL-31/IL-31RA axis using two complementary mouse models of allergic asthma. Our Th1/Th2 cytokine intratracheal instillation studies using wild-type and IL-31RA knockout mice revealed an uncoupling of AHR from other airway pathologies typically underlying asthma, including inflammation and goblet cell hyperplasia. Consistent with the important role of IL-31RA but not IL-31, only the receptor for IL-31 was elevated in wild-type mice exposed to allergens, including HDM and SEA, and administration of IL-31 had 1) limited or no effect on MCh-induced AHR, inflammation, and goblet cell hyperplasia and 2) no significant changes in the contraction of airways in the PCLS or collagen gel-embedded ASMCs.

Our findings suggest that the expression of IL-31RA, at least in the setting of primordial knockout, is not essential for both Th1 and Th2 cytokine-induced inflammation and goblet cell hyperplasia. However, this apparently limited role of IL-31RA is consistent with a recently published study in which neutralization of IL-31 had no effect on airway inflammation in wild-type mice sensitized and challenged with ovalbumin (*59*). Moreover, a study by Neuper et al. demonstrated an elevated expression of IL-31RA but not IL-31 in the lungs of mice challenged with the timothy grass pollen allergen (*37*). The role of IL-31RA has been extensively investigated in other disease models, including pulmonary fibrosis and systemic sclerosis. In support of our findings, the loss of IL-31RA had a limited effect on the expression of Th2 cytokines and inflammation but resulted in significant improvement in lung function and pulmonary fibrosis (*25, 37, 60*). Upon repetitive injury with bleomycin, mice deficient in IL-31RA were protected from worsening lung function compared with wild-type mice (37). However, studies using SEA-induced pulmonary injury model demonstrated elevated granulomatous inflammation and Th2 responses in the absence of IL-31RA (*46*). Similarly, studies using the gastrointestinal helminth Trichuris muris infection model support elevated Th2 responses and accelerated expulsion of Trichuris with significantly decreased worm burdens in the absence of IL-31RA compared with their wild-type counterparts (*46, 59, 61, 62*). Therefore, we used an alternative mouse model of allergic asthma, using SEA as an allergen, to assess AHR, airway inflammation, goblet cell hyperplasia, and Th2 responses. Consistent with our findings in the HDM allergen model, we observed no significant differences in SEA-driven Th2 responses between wild-type and IL-31RA deficient mice but observed a significant improvement in AHR. Further, our in vivo studies using intratracheal instillation of IFN-γ and IL-13 suggest that IL-31RA-driven AHR could be downstream of both Th1 and Th2 cytokines. The lack of consistency in Th2 responses in the absence of IL-31RA could be due to the disease model used, animal housing conditions, or differences in the generation of IL-31RA knockout mice (*37, 46, 61, 62*). Nevertheless, future studies are needed to assess the effect of smooth muscle cell-specific IL-31RA deficiency on AHR and inflammation using both helminth infection and allergic asthma models. In the present study, our findings demonstrated that the loss of IL-31RA is sufficient to attenuate both IFN-γ – and IL-4/IL-13-induced AHR without inhibiting the infiltration of inflammatory cells in the lung. However, mechanistic studies are warranted to determine the molecular pathways through which IL-31RA alters the contractile functions of ASMC and AHR in asthma. These studies should be complemented by the assessment of the potential role of both soluble and full-length isoforms of IL-31RA in augmenting the contractile function of ASMC and AHR in asthma (*52*).

IFN-γ is a potent Th1 cytokine primarily produced by Th1 cells to induce AHR in asthma (*12, 63, 64*). Patients with severe asthma who respond poorly to corticosteroid therapy have been shown to have dominant Th1 responses, including elevated IFN-γ in their lungs (*12, 65*). In vivo studies using animal models of severe asthma have suggested that IFN-γ, but not IL -17, is responsible for heightened AHR (*12*). In another study, the blockade of IFN-γ attenuated Th1-cell-induced AHR, and this improvement in AHR appeared to be independent of neutrophils (*63*). Similarly, multiple preclinical and clinical studies using antibody-mediated blockade of IL-4/IL-13 signaling have demonstrated the pathogenic role of Th2 cytokines in the development of AHR in allergic asthma (*18, 50, 66*). However, the mechanisms underlying Th1/Th2 cytokine-driven AHR remain unclear. Although the hypothesis tested in this study was straightforward, it was surprising that IL-31RA, but not the Th2 T cell-derived cytokine, IL-31 was responsible for inducing AHR in allergic asthma. Nonetheless, we demonstrated that both Th1 and Th2 cytokines upregulated IL-31RA in ASMC and in the two complementary mouse models of allergic asthma. Our findings on the effect of IFN-γ -induced IL-31RA expression are supported by previous studies in which IFN-γ was shown to induce the expression of IL-31RA in dendritic cells, lung epithelial cells, and fibroblasts (_26, 31, 67, 68_). A recent study by Kobayashi et al. also demonstrated the role of IFN-γ in the induction of AHR without significantly affecting airway inflammation using intranasal administration of IFN-γ in mice (*53*). However, future studies are required to determine mechanisms that uncouple AHR from inflammation and goblet cell hyperplasia in asthma.

We also provide a novel finding that IL-31RA functions as a positive regulator of CHRM3-driven calcium signaling and contractility of ASMC. This is the first study, to our knowledge to show that the IL-31RA-CHRM3 axis induces the phosphorylation of MLC in ASMC. CHRM3 is a key muscarinic receptor expressed by smooth muscle cells in the airways and plays a major role in the contractility of smooth muscle cells in response to muscarinic ligands such as acetylcholine (*56, 58*). CHRM3 is a member of class A GPCR that can mediate ASMC contraction through both calcium-dependent and calcium-independent mechanisms (*55, 58*). The calcium-dependent smooth muscle cell contractility is dependent on the activation of a subunit of Gq and the release of inositol 1,4,5-triphosphate by phospholipase C (*69*). Our studies using both the loss of function and gain-of-function studies suggest that IL-31RA augments CHRM3-dependnet calcium release and phosphorylation of MLC. Notably, we observed a possible physical interaction between IL-31RA and CHRM3 that may contribute to enhanced calcium signaling and contractility of ASMC through stabilizing CHRM3 protein. It is possible other mechanisms that involve stabilizing CHRM3 or other proteins in macromolecular complexes that involved in calcium-dependent and calcium-independent ASMC contraction (55, 58). How these observations including the physical interaction between IL-31RA and CHRM3 are linked to elevated CHRM3-driven signaling remains unknown. Further studies are required to determine the validity of the hypothesis that IL-31RA stabilizes CHRM3 to augment intracellular calcium signaling in the contraction of ASMC and whether it extends to other CHRMs. Nevertheless, given the robust increases in CHRM3-driven calcium signaling, and ASMC contraction, it is easy to envision the scope of the IL-31RA-CHRM3 axis in inducing AHR in asthma. Our findings provide evidence of the potential therapeutic benefits of targeting IL-31RA, which is downstream of both Th1 and Th2 cytokines, to attenuate AHR. This hypothesis is important for further investigation because therapeutic antibodies targeting IL-31RA are currently in multiple phase II clinical trials for atopic dermatitis (*70*). Given these findings, it is important to determine whether neutralizing antibodies or small-molecule inhibitors of IL-31RA can alter muscarinic signaling in ASMC. If so, therapeutic efficacy might be improved with these antibodies and new inhibitors, which may reduce the expression of IL-31RA and/or muscarinic signaling in ASMC. Our findings suggest that highly effective anti-AHR therapies are required to inhibit IL-31RA-driven contraction of ASMC, and it may need to be combined with therapeutics that mitigate inflammation and goblet cell hyperplasia.

In summary, we have described the role of IL-31RA in the induction of AHR, with limited effects on airway inflammation and goblet cell hyperplasia, and this role is independent of its ligand IL-31. Importantly, we identified a novel mechanism underlying ASMC contractility that involve IL-31RA dependent increases in CHRM3-driven calcium signaling and phosphorylation of MLC. Together, these results suggest an important role for IL-31RA in the regulation of AHR and identifying the molecular events underlying IL-31RA-driven AHR may lead to the development of novel therapeutic approaches against AHR in allergic asthma.

## MATERIALS AND METHODS

### Animals

Wild-type and interleukin receptor IL-31RA^-/-^ mice with a C57BL/6 background were used for in vivo experiments. Age-matched male and female mice of 12–16 weeks of age were used for experiments with at least two repeats. Wild-type mice were obtained from the Jackson Laboratory and maintained in animal facilities. IL-31RA^-/-^ mice used in this study were previously described (*26, 37, 59*). A functional loss of IL-31RA signaling has been induced in these mice through targeted deletion of 4–6 exons in the mouse genome, which encodes the cytokine-binding domain 2 of IL-31RA. All mice were housed under specific pathogen-free conditions at the Cincinnati Children’s Hospital Medical Center, a medical facility approved by American Association for the Accreditation of Laboratory Animal Care. All experimental procedures were approved by Animal Care and Use Committee of Cincinnati Children’s Hospital Medical Center.

### HDM- and SEA-induced allergic asthma mouse models

Mice were challenged with HDM extracts (*Dermatophagoides pteronyssinus*, Greer laboratories InC), a clinically important and commonly used allergen to induce asthma without the need for an adjuvant. Mice were sensitized twice with 200 µg of HDM in 200 µL of phosphate-buffered saline (PBS) through intraperitoneal injections on days 0 and 7. Mice were further challenged on days 14 and 16 through intratracheal instillation of 50 µg of HDM in 50 µL PBS. Mice were anesthetized with ketamine and xylazine to administer the allergen to the airways. Twenty-four hours following the final challenge with HDM, mice were deeply anesthetized to assess the AHR using Flexivent, and tissues were collected for biochemical and histological analyses. Control mice were sensitized with HDM and challenged with PBS.

The SEA prepared from *Schistosoma mansoni* helminth was used in other experiments to induce AHR and inflammation in mice. SEA has been used for decades as an antigen or allergen owing to its robust induction of type 2 immune responses in animal models. Previous studies have described SEA-induced asthma model (*28, 38*). Briefly, mice were sensitized with 10 µg of SEA in 200 µL by intraperitoneal injection on days 0 and 14. Two weeks later, mice were anesthetized with ketamine and xylazine, and then challenged by intratracheal instillation with 10 µg of SEA in 50 µl of PBS, twice on days 28 and 31. The day following the last challenge, the mice were euthanized and samples were collected for further analysis.

### Cytokines treatments

To evaluate the specific role of cytokines in the induction of AHR and inflammation, wild-type and IL-31RA^-/-^ mice were intratracheally administered recombinant mouse cytokines, as previously described with some modifications (*18, 20*). Mice were anesthetized by an intraperitoneal injection of ketamine/xylazine, rIL-31 (10 µg, R&D Systems), rIFN-γ (5 µg, R&D Systems), or rIL-13 (5 µg, R&D Systems) and intratracheally instilled in 50 µL of saline on days 0 and 6. Twenty-four hours after the last challenge, mice were euthanized, airway resistance was measured using Flexivent, and the lungs were collected to assess histological changes and gene expression analysis using reverse transcriptase PCR (RT-PCR). Control mice were treated with a saline solution.

### Preparation of BAL cells

Twenty-four hours after the last challenge with the allergen, the mice were anesthetized, and the lungs were lavaged with 1 mL of ice-cold and sterile PBS using a 1 mL insulin syringe through a tracheal catheter. BAL fluid was centrifuged for 5 min at 250 × g at 4 °C. Cell pellets containing BAL cells were resuspended in 500 µL of PBS. Total cell counts were determined using an automated cell counter (Thermo Fisher Scientific, Waltham, MA, USA). BAL cells were used for cytospin preparation and stained with the Diff-Quick staining kit (Thermo Fisher). Differential cell counts were determined using morphological criteria under a microscope.

### Histology

Mice were euthanized 24 h after the last allergen (HDM or SEA) challenge or after the last cytokine instillation. The lungs were inflated, fixed in 10% neutral formalin, and collected for histological analysis. Paraffin-embedded 5 µm sections of the left lobe were stained with H&E to examine associated airway inflammation and peribronchial cell infiltration. ABPAS (Alcian blue periodic acid Schiff) staining was used to assess goblet cell hyperplasia and mucus hypersecretion. Goblet cells were further counted on lung sections and expressed as the percentage of ABPAS-positive cells of the total nucleated cells of the airway epithelium, using Metamorph imaging software (Molecular Devices). At least 10 images per mouse were used to count goblet and total epithelial cells (68). Images were captured at 40x magnification using a Keyence BZ-X microscope with a focus on the airway areas to quantify the ABPAS-positive cells.

### AHR measurements

AHR was assessed in vivo using the forced oscillation technique in FlexiVent system (SCIREQ, Montreal, QC, Canada). Twenty-four hours after the last challenge, the mice were anesthetized using a ketamine/xylazine mix. Tracheotomy was performed, and the trachea was cannulated using a G20 stainless catheter and connected to the FlexiVent machine for airway resistance measurements. Increasing doses of methacholine (MCh, Sigma Aldrich), 0, 3, 6, 12.5, 25, and 50 mg/mL in PBS, were aerosolized using an ultrasonic nebulizer connected to the FlexiVent system and administrated following the manufacturer’s instructions. Lung function measurements were recorded, and airway resistances were presented as the maximum resistance (Max Rrs) values recorded for increasing MCh doses.

### PCLS model

PCLS were prepared as described previously (37, 71). Eight mice of 10 weeks of age were euthanized with a lethal dose of sodium pentobarbital. The lungs were inflated with warm 1 mL of 1.5% low melting agarose in PBS at 37 °C through a tracheal cannula, followed by 0.2 mL of air to flush the agarose. The left lung and apical and basal right lobes were used to generate PCLS. Lungs were sliced into 300 µm section in 4 °C cold HBSS without calcium using a VF-310Z vibratome (Pecisionary Instruments, Winchester, MA, USA). PCLS were collected in DMEM media without serum, and the media were changed at least four times with intermittent shaking to remove residual agarose and incubated in a humidified incubator with 5% CO_2_ at 37 °C. On the following day, the media was replaced with fresh media without serum for the same-day experiment. For cytokine treatments, PCLS were treated with IL-13 (50 ng/mL, R&D Systems) or IL-31 (500 ng/mL, R&D Systems) for 24 h in a low-serum media (1% FBS) prior to the contraction assay using increasing doses of MCh. Lung slices were allowed to rest and relax before the next dose of MCh was administered to the samples. Changes in the airway lumen area were recorded in a time-lapse setting with images captured every 5 s for 5 min following each dose of MCh using a 10x objective on a temperature-controlled Nikon inverted microscope at 37 °C with 5% CO_2_ (NIKON Ti2 widefield, Japan). Airway areas were analyzed using NIKON NIS-Elements analysis software, and changes were expressed as the percentage area of contraction compared to the initial baseline area.

### RNA isolation and real-time PCR

Lung tissues were dissociated using TRIzol (Life Technologies) using beads and a high-speed homogenizer (Thermo Fisher). RNA was isolated from lung tissues, primary mouse ASMs, or primary asthmatic human bronchial ASMC (Lonza, Walkerville,MD, USA) using an RNAeasy mini kit (QIAGEN) following the manufacturer’s instructions and as previously described in our previous studies (*28, 72*). cDNA was synthesized using Superscript III (ThermoFisher), and real-time PCR was performed using SYBR Select Master Mix (Bio-Rad) and a CFX384 Touch Real-Time PCR instrument (Bio-Rad). Data were analyzed using CFX Maestro software version 4.0. Target gene transcripts were normalized using hypoxanthine-guanine phosphor ribosyl transferase (HPRT, for murine cells and tissues) or β-actin (for human cells) as housekeeping genes. The list of primers used to measure the transcript levels of the genes of interest can be found in the supplementary material (Table S1).

### Collagen gel contraction assay

Human bronchial smooth muscle cells were obtained from Lonza (Wakersville, MD, USA), and murine ASMC were prepared via enzymatic digestion of the mouse trachea at 37 °C for 60 min, as previously described (*73*). A single-cell suspension of digested trachea was seeded on a 100 mm petri dish (n = 4/dish) using Hams/F12 culture media supplemented with 10% FBS and antibiotics. Spindle-shaped cells were allowed to grow to 80% confluence, and cells from passages 1–2 were used for the collagen gel contraction assay. ASMCs were seeded into rat tail collagen (1 mg/mL, Gibco) to collagen gel matrices embedded with ASMC, as described previously (74, 75). To determine the effect of cytokines on ASMC contraction, cells were treated with cytokines (rIL-31 500 ng/mL or rIL-13, 50 ng/mL) or media for 48 h prior to embedding them in collagen gel. Collagen gels were prepared using cells treated with cytokines and media and grown under similar conditions in Ham’s/F12 complete media. The collagen gels were detached from the walls, and images were captured at 0, 12 or 16, 24, and 48 h using a stereo microscope. The area of contraction was measured using ImageJ software and is described as the percentage of the contracted area compared with the baseline area of the gel.

### Western Blotting

Western blot analysis of smooth muscle cell lysates were performed as described earlier (72). Briefly, cell lysates were prepared using the RIPA lysis buffer containing protease and phosphatase inhibitors. After SDS–PAGE separation, proteins were transferred to nitrocellulose membrane and then were probed using specific primary antibodies CHRM3(1:1000; ab126168), phosphoMLC (1:1000; CST 3771S), totalMLC (1:1000; CST3672s), and GAPDH (1:2000 Bethyl A300-643A) followed by detection with HRP conjugated secondary antibodies. Bands were quantified using the volume integration function of the imager software, (BIORAD).

### Calcium flux assay

HEK293 cells were transiently transfected with overexpressing plasmids for IL-31RA, CHRM3 or empty control plasmids and cultured in DMEM media for 72 h (Fig S4). On day 3, cells were loaded with Fluo-4 AM (5µM) (Invitrogen) for one hour and stimulated with carbachol (10 µm) to measure changes in the fluorescence in real time for 90 sec using a Flex Station III (Molecular Devices).

### *In situ* proximity ligation assay

HEK293 cells were transiently transfected with overexpressing plasmids for IL-31RA (Origene, RC218212L1) and CHRM3 (Origene, SC119736) or control empty plasmids (pLJM1-EGFP, Addgene 19319 and pLenti-C-Myc-DDK-P2A-Puro, Origene PS100092) using Lipofectamine 3000 and cultured for 48 h. Then, cells were fixed for 5 min in ice cold methanol at 4^0^C and incubated with Duolink blocking solution (Sigma) for 1 hour at room temperature followed by overnight incubation with primary antibodies for IL-31RA (15 µg/ml, R&D Systems AF2769) and CHRM3(1:500, Abcam ab126168). The incubation was followed by three 5 min washes and incubation with PLA probes anti-goat PLUS (DUO92003) and anti-rabbit MINUS (DUO92005) for 1 h at 37^0^C as recommended in the Duolink PLA kit (Sigma). After three washes and cells were incubated with the hybridization solution containing DNA ligase for 30 min at 37°C. Cells were washed and incubated with the amplification-polymerase at 37°C for 100 min, washed three times and mounted with DAPI Mounting Medium. To identify the membrane localization of *in situ* PLA signals, the cells were counterstained with cholera toxin conjugated with Alexa Fluor 488 (1:200, Thermo Fischer C34775). Fluorescence images were acquired using a Nikon A1R confocal laser scanning microscope and analyzed using Imaris image analysis software.

### Statistical Analysis

Data were analyzed using Prism (version 9; GraphPad, San Diego, CA, USA). Quantitative data are presented as bars with individual dots, including mean ± SEM. Data were considered statistically significant when the p-values were less than 0.05. Two tailed Student’s t-test was used to compare the means between the two groups. For multiple comparisons, one-way ANOVA with post hoc Tukey’s test or 2-way ANOVA with post hoc Sidak’s test was used.

## Acknowledgments

We thank the veterinary services and core facilities at the Cincinnati Children’s Hospital Medical Center and the University of Cincinnati. We also thank Eric Smith from University of Cincinnati for editing the manuscript.

## Author contributions

Conceptualization: SKM; Methodology: SKM, DJKY, SA, SY; Investigation: SKM, DJKY, SA, SY; Visualization: DJKY, SA; Supervision: SKM, BRG, FXM; Writing-original draft: SKM, DJKY; Writing-review and editing: SKM, DJKY, SA, FXM.

