## Supplementary Data for "Interleukin 31 receptor alpha induces airway hyperresponsiveness in asthma"

**This PDF file includes:**

Figures S1, S2, S3, S4, S5 and S6

Tables S1

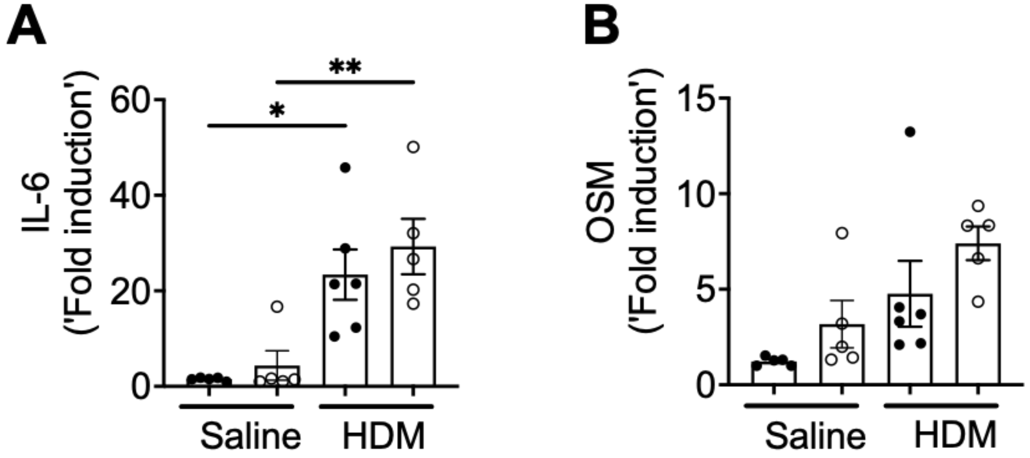


**Figure S1. The loss of IL-31RA has no effect on the expression of IL-6 and OSM in the lungs of HDM-challenged mice compared to saline-treated mice.** Quantification of IL-6 **(A)** and OSM **(B)** transcripts in the lungs of wildtype and IL-31RA$^{-/-}$ mice treated with HDM or saline.

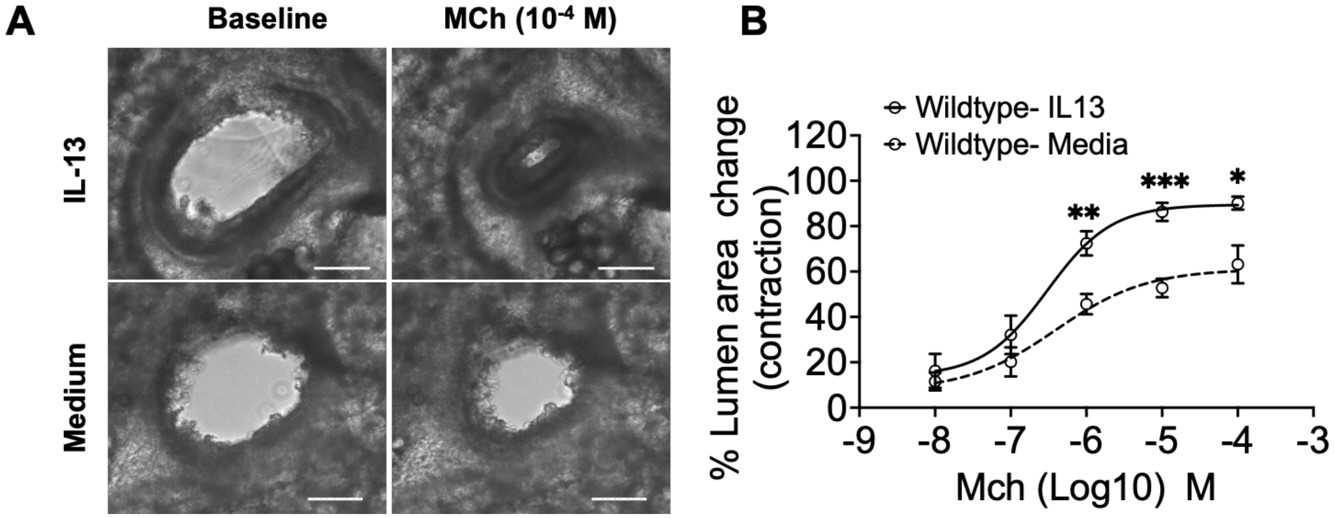


**Figure S2 IL-13 induces contraction of airways in PCLS of wild-type mice. (A)** Representative images of MCh-induced airway lumen contraction in PCLS of wild-type mice treated with IL-13 or media for 24 hrs. Images were captured at 10X magnification, scale bar 150um. **(B)** The percent of airway lumen area contraction in response to increasing doses of MCh between saline and IL-13 treated PCLS from wild-type mice. Data are shown as means ± SEM, n=5-8/group; Two-way ANOVA test with * $p<0.5$.

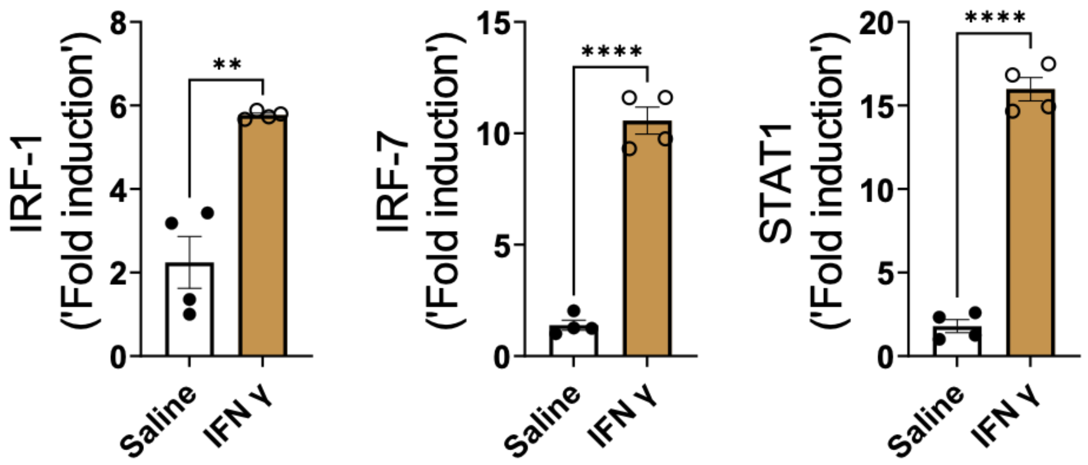


**Figure S3. Upregulation of IFNγ-induced genes in the lungs of IFNγ treated mice.** Wildtype mice were treated intratracheally with IFNγ ( 5 μg) or saline on days 0 and 6, then sacrificed 24 hours after the last challenge. To confirm the effects of IFNγ, the expression of IFNγ-specific gene transcripts including IRF1, IRF7 and STAT1 were measured in the lungs using RTPCR. Data are shown as means ± SEM, n=4/group. Two-tailed Student *t*-test with no statistical significance between groups.

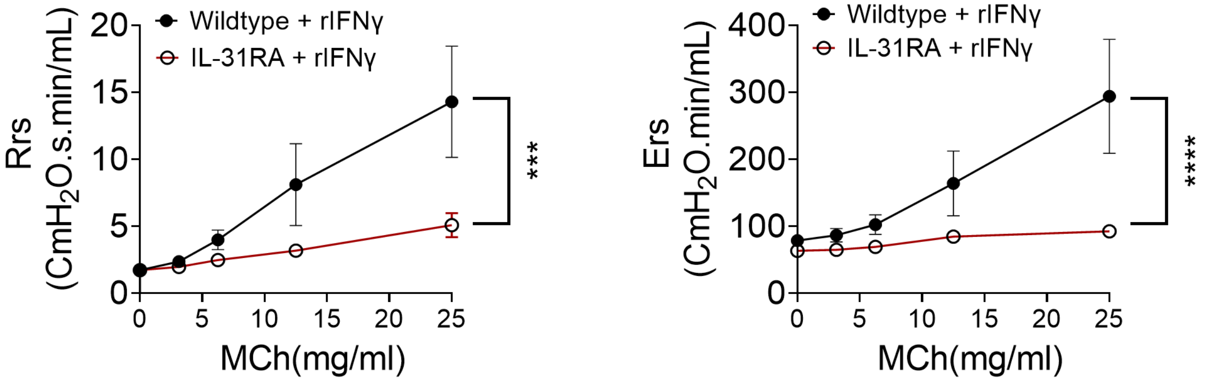


**Figure S4. IFN-γ induced responses in the wild-type and IL-31RA$^{-/-}$ mice**. Wild-type and IL-31RA$^{-/-}$ mice were treated intratracheally with IFN-γ (5 µg) on days 0 and 6. The methacholine-dependent increase in resistance (Rrs) and elastance (Ers) were measured using Flexivent on day 7. Data are shown as mean ± SEM, n = 6/group. Two-way ANOVA was used, * $P < 0.05$.

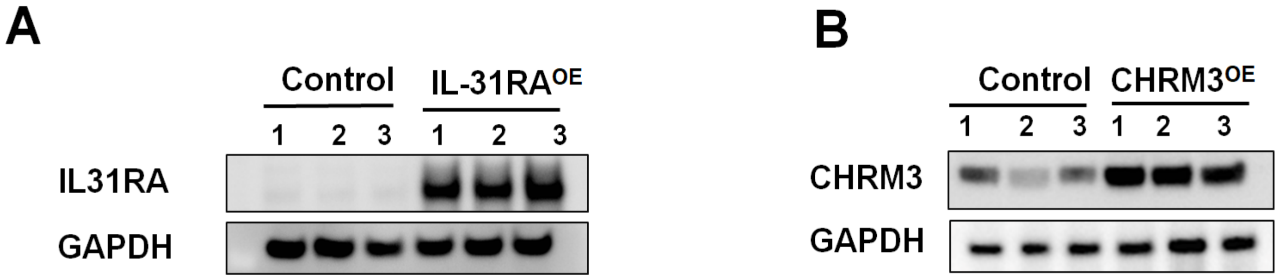


**Figure S5. Overexpression of IL-31RA and CHRM3 in HEK 293 cells. (A)** HEK293 cells were transiently transfected with overexpressing plasmid for IL-31RA for 72 h and cell lysates were immunoblotted with IL-31RA and GAPDH. **(B)** HEK293 cells were transiently transfected with overexpressing plasmid for CHRM3 for 72 h and cell lysates were immunoblotted with CHRM3 and GAPDH. Data are shown as means ± SEM, n = 3/group.

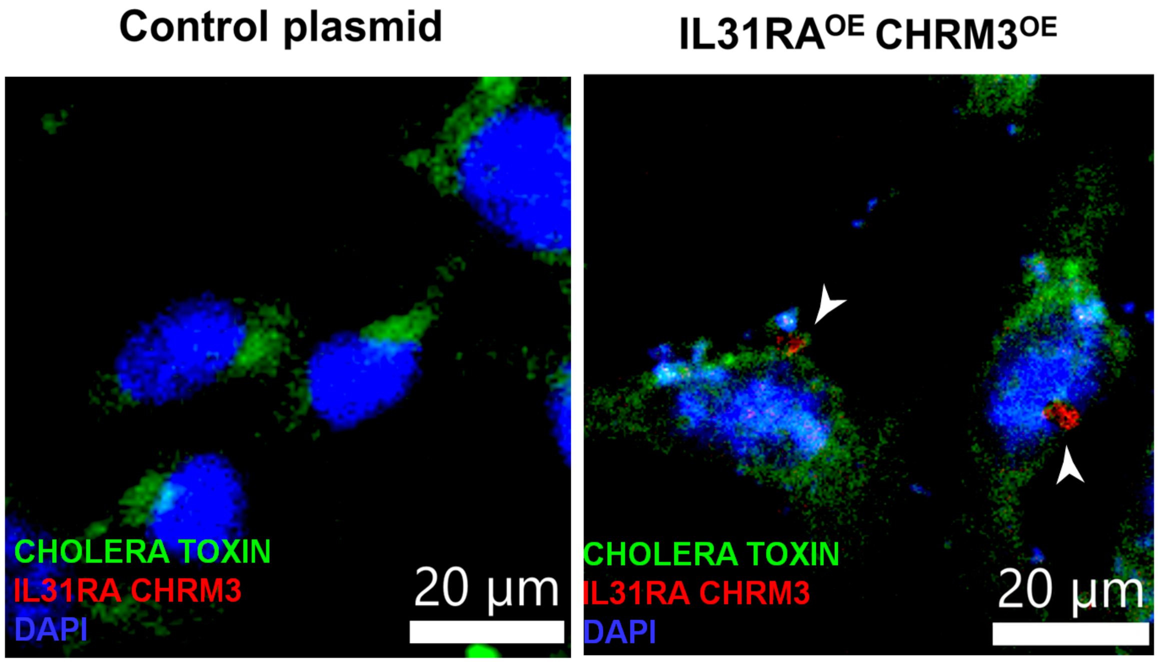


**Figure S6. Colocalization of the IL-31RA-CHRM3 complex with cholera toxin.** HEK293 cells were transiently transfected with overexpressing plasmid for control (plenti-cmyc-DDK -P2A-Puro) or plasmids overexpressing IL-31RA and CHRM3 for 48 h and stained for cholera toxin and the IL-31RA-CHRM3 complex. The IL-31RA-CHRM3 complex formation was visualized using hybridization probes labeled with Alexa 594 (Red). The plasma membrane was stained with cholera toxin subunit b conjugated with Alexa Fluor 488 and the nuclei were stained with DAPI (blue). The white arrowhead represents colocalization between the cholera toxin subunit b and puncta of the IL-31RA-CHRM3 complex. Images were captured at 40X magnification. Scale bar, 20 µm.

**Table S1. List primers used for qPCR transcript expression of genes of interest**

| Gene ID | FORWARD | REVERSE |
|---|---|---|
| **hIL-31** | GCCCAGCCGCCAAAC | GCTGTCTGATTGTCTTGAGATATGC |
| hIL-31RA | TAGTACCAGATCATCTGTGT | TTAGACTTCTCCCTTGGTGTGC |
| hβ ACTIN | CCA ACC GCG AGA AGA TGA | CCA GAG GCG TAC AGG GAT AG |
| mARG1 | GGAAAGCCAATGAAGAGCTG | GCTTCCAACTGCCAGACTGT |
| mCCL11 | AGAGCTCCACAGCGCTTCT | GCAGGAAGTTGGGATGGA |
| mCCL24 | GCAGCATCTGTCCCAAGG | GCAGCTTGGGGTCAGTACA |
| mCHIL3L | GTAGCACATCAGCTGGTAGGA | AAAGGCAAAACTGTTAGCCAAGG |
| mFIZZ1 | CCCTCCACTGTAACGAAGACTC | CACACCCAGTAGCAGTCATCC |
| mGOB5 | AGGAAAACCCCAAGCAGTG | GCACCGACGAACTTGATTTT |
| mHPRT | GCCCTTGACTATAATGAGTACTTCAGG | TTCAACTTGCGCTCATCTTAGG |
| mIFNγ | AGAGCCAGATTATCTCTTTCTACCTCAG | CCTTTTTCGCCTTGCTGTTG |
| mIL-4 | ACGAGGTCACAGGAGAAGGGA | AGCCCTACAGACGAGCTCACTC |
| mIL-5 | TGACAAGCAATGAGACGATGAGG | ACCCCCACGGACAGTTTGATTC |
| mIL-6 | GCTACCAAACTGGATATAATCAGGA | CCAGGTAGCTATGGTACTCCAGAA |
| mIL-10 | CAGAGCCACATGCTCCTAGA | GTCCAGCTGGTCCTTTGTTT |
| mIL-13 | CCTCTGACCCTTAAGGAGCTTAT | CGTTGCACAGGGGAGTCTT |
| mIL-17 | CAGGGAGAGCTTCATCTGTGT | GCTGAGCTTTGAGGGATGAT |
| mIL-31 | TCTTACCGTCGCCATGATCT | GCACCGAAGGACAAGCTG |
| mIL-31RA | CAGAATTCTCCACAGGTCCAG | TGGAGCAAAGAAGAAACCAGA |
| mIRF-1 | GTTGTTTACAGCGTGTGGCTT | AGCCAGCAAAAGACTCCCAT |

| | | |
|---|---|---|
| mIRF-7 | TCCAGTTGATCCGCATAAGGT | CTTCCCTATTTTCCGTGGCTG |
| mMUC4 | CAGGCAAGGTTGAGGTATCC | CATGCATAAGAAAAGGCGGCA |
| mMUC5AC | ACTTCAACGGCAGTCCAAAA | CTCAAGGGGTGTCAGCCTAA |
| mOSM | TGCTCCAACTCTTCCTCTCAG | CAGGTTTTGGAGGCGGATA |
| mSOCS3 | ATTTCGCTTCGGGACTAGC | AACTTGCTGTGGGTGACCAT |
| mSTAT1 | CCCCTGAAGTATCTGTACCCCA | CTCAGGCACTCACCTTCCTTT |
| mTNFα | GCCTCTTCTCATTCCTGCTTGT | GGCCATTTGGGAACTTCTCAT |
